# A molecular census of midbrain dopaminergic neurons in Parkinson’s disease

**DOI:** 10.1101/2021.06.16.448661

**Authors:** Tushar Kamath, Abdulraouf Abdulraouf, SJ Burris, Vahid Gazestani, Naeem Nadaf, Charles Vanderburg, Evan Z Macosko

**Affiliations:** Broad Institute of Harvard and MIT, Stanley Center for Psychiatric Research, 75 Ames Street. Cambridge, MA, USA; Harvard Graduate Program in Biophysics, Harvard University; Tri-Institutional MD–PhD Program, Weill Cornell Medical College, Rockefeller University and Memorial Sloan Kettering Cancer Center, New York, NY, USA; Massachusetts General Hospital, Department of Psychiatry, 55 Fruit St. Boston, MA, USA

## Abstract

Midbrain dopamine (DA) neurons in the substantia nigra pars compacta (SNpc) project widely throughout the central nervous system, playing critical roles in voluntary movements, reward processing, and working memory. Many of these neurons are highly sensitive to neurodegeneration in Parkinson’s Disease (PD), and their loss correlates strongly with the pathognomonic symptoms. To characterize these populations molecularly, we developed a protocol to enrich and transcriptionally profile DA neuron nuclei from postmortem human SNpc of both PD patients and matched controls. We identified a total of ten distinct populations, including one that was primate-specific. A single subtype, marked by the gene *AGTR1*, was highly susceptible to degeneration, and was enriched for expression of genes associated with PD in genetic studies, suggesting many risk loci act within this subtype to influence its neurodegeneration. The *AGTR1* subtype also showed the strongest upregulation of *TP53* and its downstream targets, nominating a potential pathway of degeneration *in vivo*. The transcriptional characterization of differentially disease-vulnerable DA neurons in the SNpc will inform the development of laboratory models, enable the nomination of novel disease biomarkers, and guide further studies of pathogenic disease mechanisms.

## Introduction

The degeneration of midbrain dopaminergic (DA) neurons within the pars compacta region of the substantia nigra (SNpc) is a pathological hallmark of Parkinson’s disease (PD) and Lewy body dementia (LBD)^1^. Nonetheless, some SNpc neurons are observed to survive even into late stages of disease^2–7^, suggesting differential vulnerability to degeneration. A clearer understanding of the specific molecular characteristics of vulnerable neurons, and the associated cascade of molecular events that lead to their demise, could provide opportunities to refine laboratory models of PD, and aid in the development of disease-modifying or cell-type-specific therapies^8^.

Recent advances in single-cell RNA-sequencing (scRNA-seq) technology^9–11^ and its application to the human postmortem brain, have begun to reveal cell-type-specific changes in several brain diseases^12–16^. Some recent studies have profiled human SNpc^17–19^, but the proportion of DA neurons is extremely small--considerably lower than in murine SNpc--making it challenging to accumulate sufficient numbers of these profiles to robustly compare across subjects. Enrichment is especially necessary to recover DA neurons from PD and LBD tissues, which show an even greater dropout of these cells.

## A map of human midbrain dopaminergic neurons

We developed a protocol, based on fluorescence-activated nuclei sorting (FANS), to enrich midbrain DA neuron nuclei for use in single-nucleus RNA-seq (snRNA-seq) (Fig. 1A, Methods). In a scRNA-seq dataset of mouse SNpc^20^, we identified the transcription factor *Nr4a2* as specific (AUC = 0.76, Extended Data Fig. 1A, Methods) to mammalian midbrain DA neurons. We isolated nuclei from eight neurotypical donors (Extended Data Table 1), and performed snRNA-seq on both NR4A2-positive and NR4A2-negative nuclei (Extended Data Fig. 1B, Methods), to comprehensively profile all cell types in the SNpc. In total, we profiled 184,673 high-quality nuclei, 43.6% of which were from the NR4A2-positive cytometry gate (Methods, Fig. 1B). The NR4A2-sorted profiles were 70-fold enriched for DA neurons (Fig. 1B, Extended Data Fig. 1H,K), providing a total of 15,684 DA profiles. We performed clustering analysis of each donor to assign non-DA profiles to one of seven other main cell classes (Methods, Extended Data Fig. 1G-L). Across all samples, we profiled nuclei deeply (median number of unique molecular identifiers, UMIs, per individual = 8810, median number of genes per individual = 3590, Extended Data Fig. 1C-D) and consistently across major cell types, with gene and transcript counts matching well with expected variation per cell type (Extended Data Fig. 1E-F).

**Figure 1:**
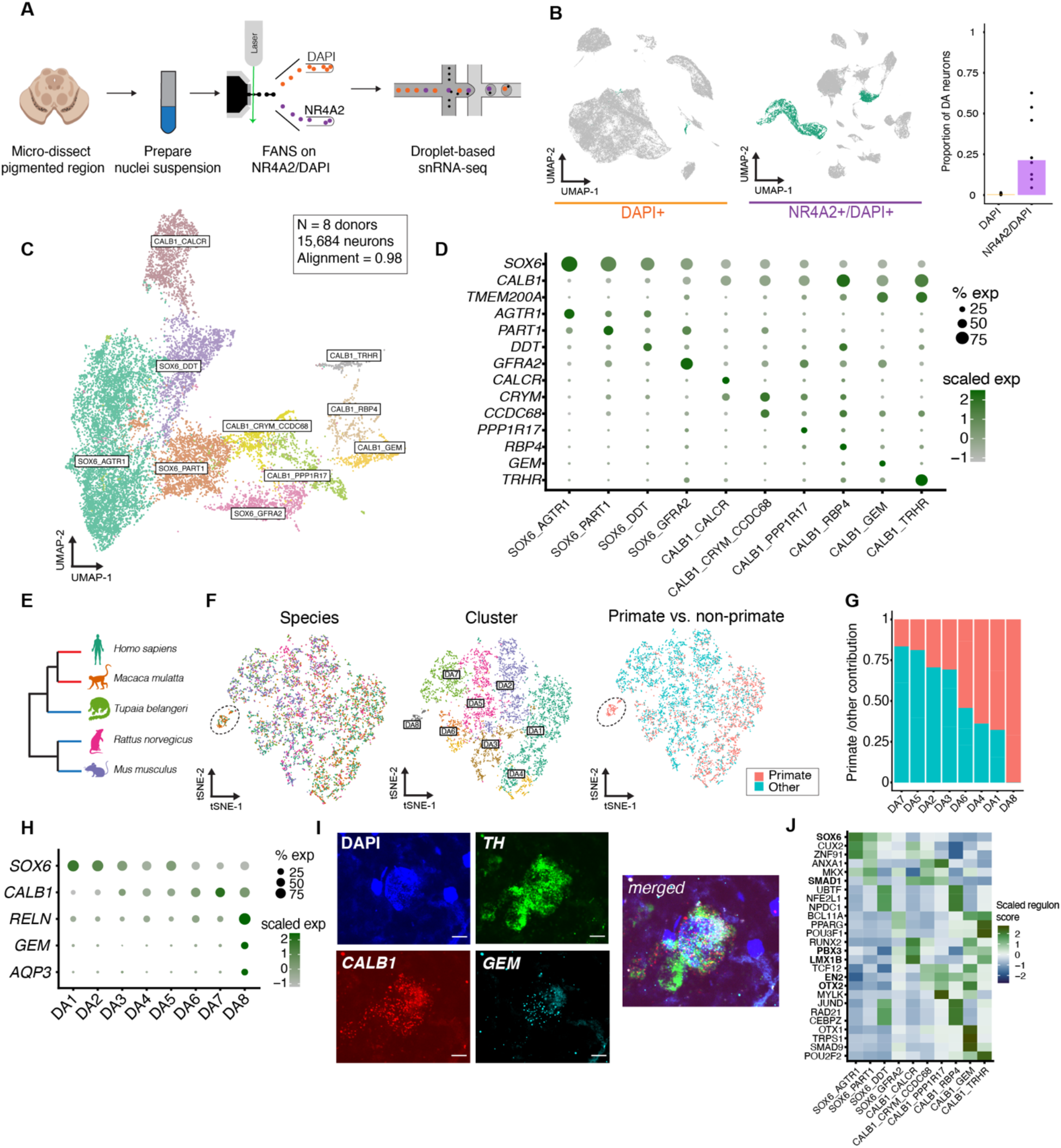
A molecular census of DA neurons in the human substantia nigra. **a,** NR4A2 antibody-based enrichment strategy and snRNA-seq profiling workflow. **b,** UMAP representation of 104,097 NR4A2- (left) and 80,576 NR4A2+ (middle) nuclei. Profiles colored green are from clusters identified as DA neurons. Right, bar plot showing the proportion of DA neurons per individual for NR4A2-versus NR4A2+ libraries (median fold enrichment = 70). **c**, UMAP representation of 15,684 DA neuron nuclei colored by cell type. **d,** dot plot showing expression of selected marker genes across the DA clusters. **e,** Dendrogram showing phylogenetic relationships amongst the five species samples in this study. Red branches denote primate species; non-primate are denoted by blue. **f**, UMAP representation of 5,904 DA neuron nuclei (2,481 primate cells and 3,423 non-primate cells) colored by species (left), cluster identity (middle), and primate versus not primate (right). **g,** Stacked bar plot showing the proportion of primate profiles (red bars) in each cluster. **h**, Dot plot of genes that mark the primate-specific cluster DA8 shown in **f. i**, Representative image of single molecule fluorescence *in situ* hybridization showing co-localization of *TH* (green), *CALB1* (red), and *GEM* (cyan) in the human SNpc. **j,** Heatmap of regulon activity from SCENIC (Methods) scaled across all 10 DA neuron subtypes of top 3 differentially expressed regulons per DA subtype. Bold indicates those transcription factors previously identified as important for midbrain DA differentiation.

To identify subtypes of DA neurons, we performed LIGER^17^ on the 15,684 DA profiles, identifying 10 transcriptionally-distinct subpopulations (Methods, Fig. 1C) with minimal inter-individual variation in our analysis (see Methods, alignment score = 0.98, Extended Data Fig. 1M). Four DA clusters expressed *SOX6*, while the other six expressed *CALB1*, recapitulating a well-defined developmental axis of variation within midbrain DA neurons^21^ (Extended Data Fig. 1N). Further, the proportions of these broad subtypes matched well with stereotactic estimates of the SNpc^22^, suggesting no intrinsic bias in our tissue sampling method (Extended Data Fig. 1O). Within the more diverse *CALB1* group, we identified one particularly transcriptionally distinct set of subtypes marked by the expression of *TMEM200A* and *RELN* (Fig. 1C).

## A primate-specific dopaminergic cell type

Although recent profiling studies of primates have shown strong evolutionary conservation of cell types in mice, some primate-specific specializations have been reported^23,24^. To investigate the evolutionary conservation of our 10 human DA populations, we sampled profiles from five species spanning three phylogenetic orders: *Primate, Scandentia*, and *Rodentia* (Fig. 1E). The integrative analysis (Methods) identified eight clusters (Fig. 1F) that recapitulated much of the diversity from the human analysis (Extended Data Fig. 2A-B).

Examining the species contributions to each cluster, we found that one, marked by expression of *CALB1, FAM83B*, and *GEM*, among other markers, (Fig. 1H, Extended Data Fig. 2C), was solely composed of profiles derived from the primate brain (Fig. 1F). Two-way alignments between human/mouse and human/macaque further confirmed the primate specificity of this population (Extended Data Fig. 2D-E). Using single-molecule fluorescence *in situ* hybridization^25^ (smFISH), we identified DA cells expressing *CALB1* and *GEM* in the human SNpc (Fig. 1I, Extended Data Fig. 2G) but not in the mouse brain (Extended Data Fig. 2F). The primate SNpc is known to have an expanded dorsal tier population and is enriched for neurons that express *CALB1*^26,27^. As such, we hypothesize that these primate-specific cells may reside in the dorsal tier of the SNpc and could be one of many neurons in this region that make atypical projections to cortex, as has been previously observed in primate brain^26,27^.

## Analysis of regulatory elements across dopaminergic neurons

The identification of regulatory elements that drive the molecular identity of DA neurons can inform differentiation protocols for *in vitro* studies of dopamine neurons in PD as well as drive the refinement of cellular replacement therapies for PD^28^. To understand the regulatory networks that may drive such transcriptional variation, we used SCENIC (Single-Cell rEgulatory Network Inference and Clustering^29^, to identify 84 regulons (Extended Data Fig. 3) that were significantly active (Wilcoxon rank-sum test p < 0.05 and area under the curve (AUC) > 0.7, Methods) in these 10 major cell populations. Indeed, the top three transcription factors (TFs) ranked by their AUC per DA subtype contained a number of TFs that specify DA neurons^30^, including *SOX6, OTX2, SMAD1, PBX3, LMX1B*, and *EN2* (Fig. 1J). Even within the more homogeneous *SOX6* axis, we identified several TFs with differential activity across subtypes, including *SMAD1*^31^, as well as TFs not previously implicated in DA neuron differentiation, such as *CUX2, ZNF91, UBTF*, and *NFE2L1*.

## Nominating compositional cell type changes in Parkinson’s disease

To identify the molecular and cellular alterations in association with PD, we sampled an additional 202,810 high-quality nuclei (median number of genes per individual = 3108, median number of UMIs per individual = 7177, Extended Data Fig. 4A-B) using our sorting procedure from 10 age-matched and postmortem interval-matched (Extended Data Fig. 4C,D, and H) individuals with documented pathological midbrain DA neuron loss, and a clinical diagnosis of either PD or Lewy body dementia (LBD) (Extended Data Table 2). Integrative analysis of these donors (Methods) identified 68 transcriptionally-defined subpopulations (Extended Data Fig. 4I-N) from our eight major cell classes (Extended Data Fig. 4E-F), with minimal batch-dependent variation (alignment scores: 0.61-1, median alignment score across cell types = 0.76, Extended Data Fig. 4G).

We assessed the differential abundance between PD and aged control samples of both the major cell classes and the molecularly-defined subtypes. Among major cell classes, DA neurons showed the largest significant (p < 0.05, Wilcoxon rank-sum test) decline as a fraction of cells per individual (Extended Data Fig. 5A). To assess differential abundance across all 68 molecularly-defined subpopulations, we employed MASC (Mixed effect Association of Single Cells), a tool that nominates proportional alterations while controlling for batch variation and individual-level covariates that influence cell proportions^32^. In total, we detected 11 subpopulations that were significantly differentially altered (FDR-adjusted p-value < 0.05, absolute(log_2_ Odds Ratio) > 0) in association with PD/LBD (Fig. 2B). One proportionally-increased population was a subset of microglia expressing *GPNMB* (Extended Data Fig. 5B), which has been identified as a marker of disease-associated microglia in transcriptomic studies of Alzheimer’s disease (AD)^33,34^. We also identified the induction of a *VIM/LHX2*+ astrocyte subtype (Extended Data Fig. 5C) whose reactive markers suggest roles in responding to the degenerative changes in PD/LBD SNpc.

**Figure 2:**
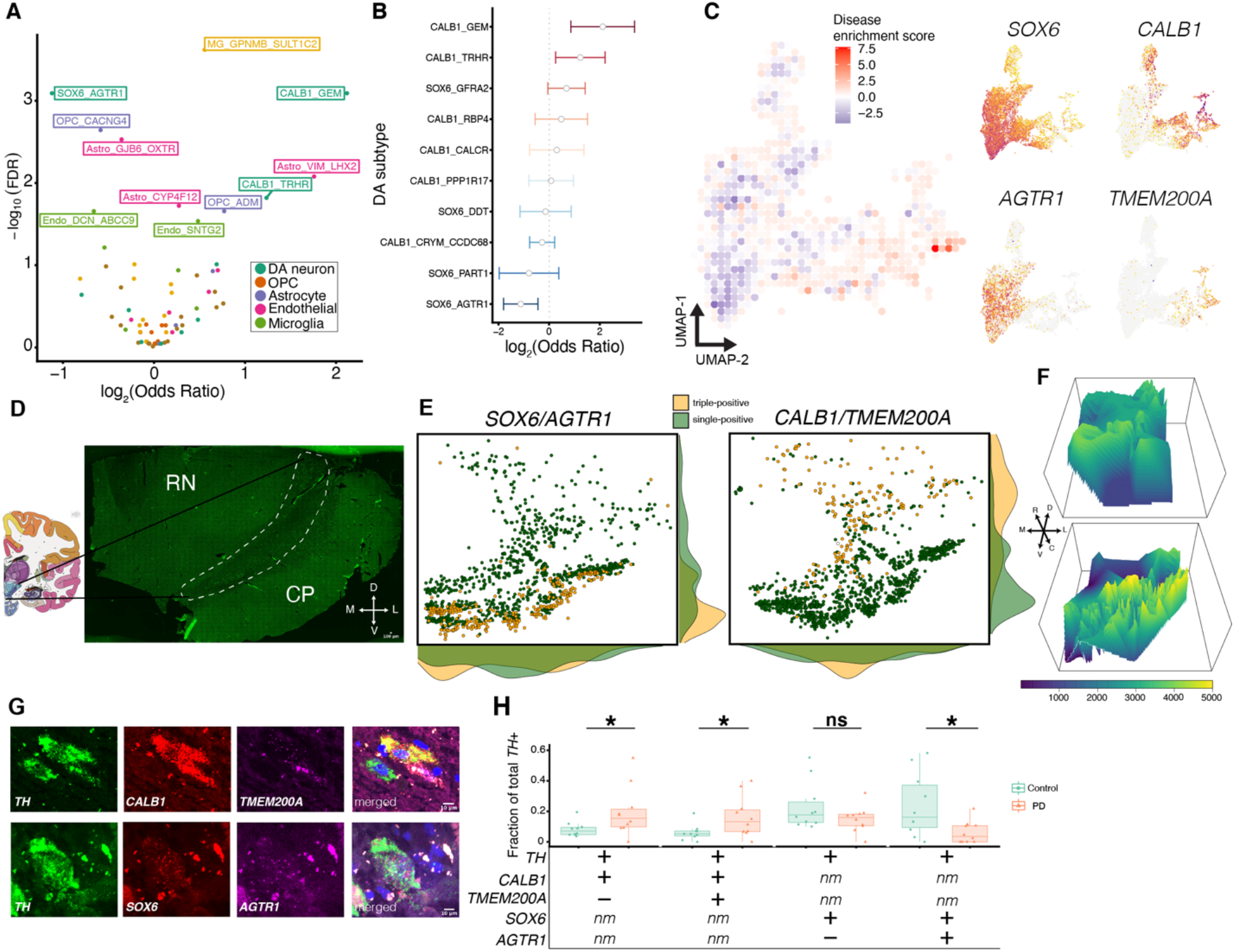
Quantification of DA subtype vulnerability to PD-associated degeneration. **a,** Volcano plot showing the odds ratio and FDR-adjusted p-value computed by MASC for each of the 68 clusters identified in the SNpc snRNA-seq analysis. The labeled clusters are those that are significantly (FDR-adjusted p-value < 0.05) increased or depleted in association with PD/LBD. **b,** Odds ratio estimate of 10 dopaminergic subpopulations as identified by MASC. Width of bar corresponds to 2.5 times the standard deviation estimates from MASC. Bars that cross zero (dotted line) are not statistically significant (FDR-adjusted p-value < 0.05). **c,** Left: Disease enrichment score (see Methods) overlaid onto a binned UMAP representation of an integrative analysis of both PD/LBD and control DA neurons. Right: Expression of select genes used to validate subtype vulnerability are plotted on UMAP representation of DA neurons. **d,** Allen human brain atlas coronal section showing the anatomical location of SNpc. Inset: representative image of tiled midbrain from one individual. Dotted outline indicates area occupied by *TH*+ cells, SNpc. RN = red nucleus, CP = cerebral peduncles. Arrows indicate orientation, D = dorsal, L = lateral, M = medial, V = ventral. **e,** Representative plates of stereotactic localization of two subpopulations in the midbrain. Left = triple-positives in yellow are *TH+/SOX6+/AGTR1*+ cells, right = triple-positives in yellow are *TH+/CALB1+/TMEM200A*+ (Methods). Margin plots are densities of the single- or triple-positive neurons. **f,** Threedimensional reconstruction of *CALB1+/TMEM200A*+ (top) and *SOX6+/AGTR1*+ (bottom) DA subtypes. All six slides were registered and cell numbers were interpolated to create the surface plots shown (Methods). Shading represents the interpolated number of neurons for each subtype. **g**, Representative images of triple positive cells one for disease-resistant (top, *TH+/CALB1+/TMEM200A*+) and disease-vulnerable (bottom, *TH+/AGTR1+/SOX6*+) DA populations. Scale bar = 10 μm. **h**, Box plot of proportions of four DA populations across 10 PD and 10 control SNpc tissue donors, determined by counting smFISH images from the two staining procedures (3,258 DA neurons counted for first assay and 2,081 DA neurons for second assay) described in **g**. For boxplots, median is center line, and interquartile range is shaded according to disease status. All dots represent an individual case for each subtype as a fraction of total *TH*+ cells counted. +, positive for marker; −, negative for marker; nm, not measured; *, p < 0.05 (Wilcoxon rank-sum test, see Methods); ns, not significant.

The cluster with the largest significant depletion was a DA subtype marked by *SOX6* and *AGTR1*. A downsampling analysis (Extended Data Fig. 5D) across cells and replicates confirmed that this DA population is consistently the most significantly depleted. Conversely, the cluster with the largest proportional increase was a distinct DA population marked by the expression of *CALB1* and *GEM*. An additional *TRHR*+ cluster that, together with the *GEM*+ population, expresses *TMEM200A*, was also proportionally increased in disease samples (Fig. 2C). We developed a metric to visualize disease-associated enrichment or depletion within the low-dimensional embedding of the jointly analyzed cell profiles, identifying a gradient of susceptibility (Fig. 2C, Methods) that correlated well with the expression of *AGTR1* and *TMEM200A* (Fig. 2C).

## Localization and validation of resistant and vulnerable dopaminergic neurons

It has been observed that PD-associated degeneration follows a spatial gradient, wherein the dorsal tier of the pars compacta exhibits far less DA neuron loss than the ventral tier ^6,35,36^. To localize our altered DA neuron populations, we performed smFISH across the full dorsal-ventral, medial-lateral, and rostral-caudal axes of one postmortem control human midbrain (Fig. 2D, Extended Data Fig. 6A-B). We found the *SOX6+/AGTR1*+ cell population, the most susceptible cell type, localizes to the ventral tier of the SNpc (Fig. 2E) while the most resistant population (*CALB1+/TMEM200A*+) was heavily enriched in dorsal tier (Fig. 2E). Across the rostral-caudal axis, these populations were consistently present throughout the midbrain, though at variable total levels (Fig. 2F, Extended Data Fig. 6A), suggesting the patterns of PD-associated degeneration are organized as a reticulum, rather than clearly demarcated pockets^37,38^.

Our flow cytometry procedure to isolate DA nuclei relies on protein expression of *NR4A2*, which has shown to be downregulated in DA neurons in PD^39^. To address these and other potential confounders, we performed smFISH on an additional 10 postmortem frozen midbrains from neurologically normal individuals and 10 from individuals with PD, matched for age and postmortem interval (Extended Data Table 3, Extended Data Fig. 6C-D). We assayed the proportional representations of four neuron subtypes (Fig. 2G) and counted a total of 5,339 individual DA neurons (Extended Data Fig. 6E) across 40 full SNpc sections. We confirmed that the *CALB1+/TMEM200A*+ DA cells were selectively enriched in PD (Fig. 2H, Wilcoxon ranksum test p < 0.05), as were the *CALB1+/TMEM200A-* cells, though the log-fold change difference was lower (Fig. 2H, log_2_ fold-change = 1.13 for the *TMEM200A-* versus log_2_ foldchange (FC) = 1.32 for the *TMEM200A*+ group, Supplementary Table 1), supporting the hypothesis that the *TMEM200A*+ populations are selectively resistant to PD-associated degeneration. Finally, we confirmed the selective depletion of the *SOX6+/AGTR1*+ dopamine subpopulation (Wilcoxon rank-sum test p < 0.05, log_2_FC = −2.1, Supplementary Table 1) relative to the *SOX6+/AGTR1-* population, whose depletion was not statistically significant (Wilcoxon rank-sum test p > 0.05). These results corroborate our snRNA-seq analysis, nominating *AGTR1* as a marker for the DA neuron subtype that is the most sensitive to PD/LBD-associated degeneration.

## Enrichment of heritable risk for Parkinson’s disease in the degenerating dopaminergic subtype

Genetic association studies for PD have identified both rare and common risk variants, but whether they directly influence neurodegeneration within DA neurons, or through other resident cell types, remains unclear. We examined the relative enrichment of genes ranked by their association with common variants in PD with markers of our SNpc cell classes^40,41^ (Methods), as well as markers of cell classes we defined by additional profiling of 46,872 nuclei from human dorsal striatum of four neuropathologically-normal individuals (Extended Data Fig. 7A-F; Extended Data Table 4). We observed strong, significant enrichment (Bonferroni-corrected p-value < 0.05) for PD-associated genes (Methods, Extended Data Fig. 8A) within DA neurons, in agreement with a recent analysis of mouse single-cell datasets^42^. In contrast, significant enrichment (Bonferroni-corrected p-value < 0.05) of genes associated by common variant studies of Alzheimer’s disease was identified only in microglia/macrophage cluster markers in the SNpc and striatum (Fig. 3A)^42,43^. Next, we tested for enrichment of disease risk gene expression across the 68 transcriptionally defined subpopulations (Methods). We found the largest and only statistically significant (Bonferroni corrected p-value < 0.05, Methods) enrichment of PD genetic risk genes within the *SOX6+/AGTR1*+ cell subtype (Fig. 3B).

**Figure 3:**
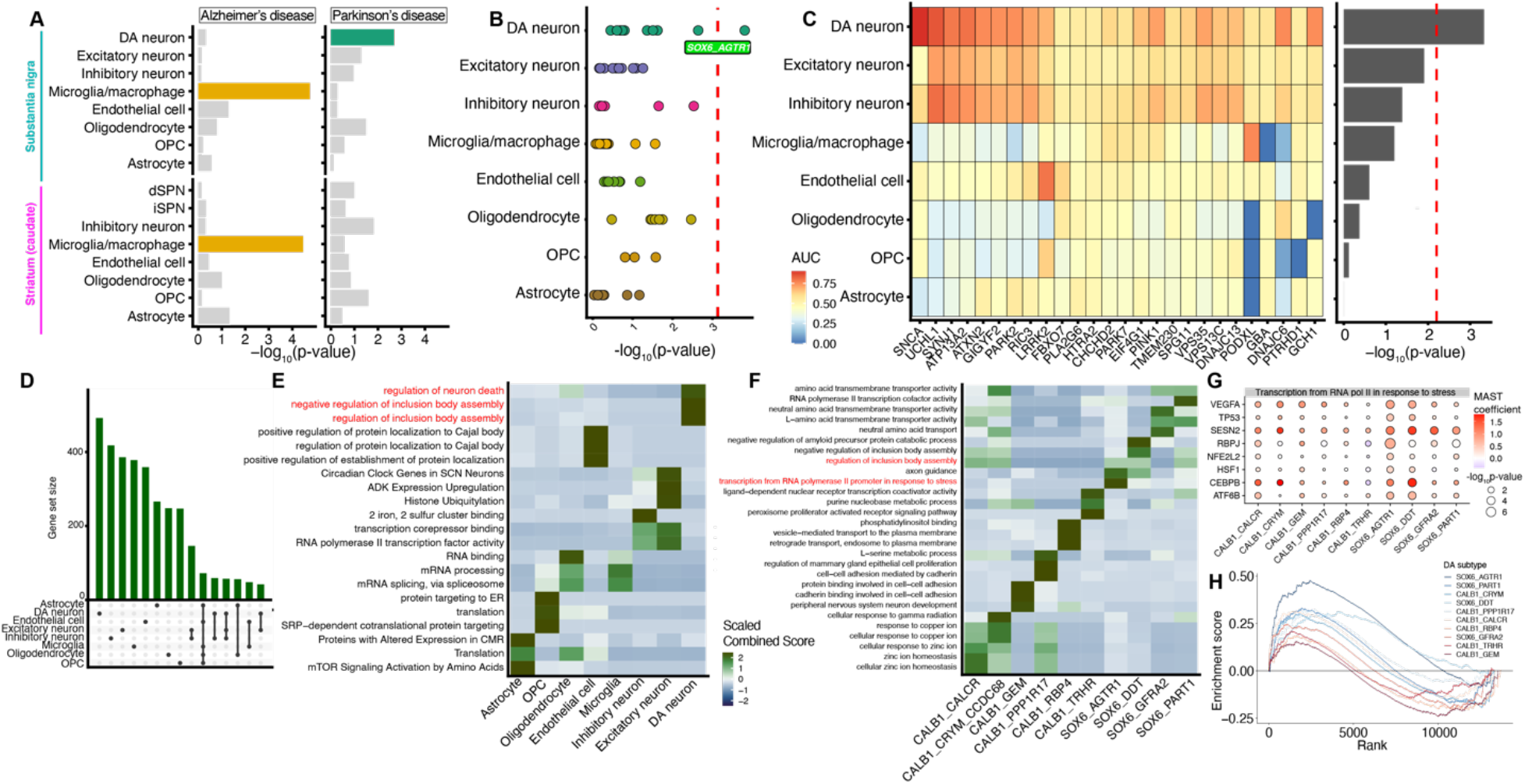
Genetic enrichments and differential expression within DA subtypes. **a,** Bar plot of −log_10_-transformed p-values from MAGMA enrichment of Alzheimer’s (left) and Parkinson’s disease (right) across 16 cell types from dorsal striatum (caudate) and SNpc. Bars are colored for significantly (Bonferroni-corrected p-value < 0.05) enriched cell types. **b**, Dot plot of −log_10_-transformed p-values for MAGMA analysis of PD genetic risk in the 68 transcriptionally defined SNpc clusters. Clusters are grouped on the y-axis by major cell class. Red dotted line indicates Bonferroni significance threshold (p < 0.05). **c,** Left: Heatmap of expression of 26 familial PD genes^44^, colored by area under the curve (AUC) statistic (Presto, see Methods). Right: Bar plot of −log_10_-transformed p-values from a Fisher’s exact test comparing the overlap between markers for major cell types (see Methods) and familial PD genes. **d**, Upset plot of top 1000 upregulated differentially expressed genes (Methods) across eight major cell classes. **e,** Heatmap of Combined Score obtained from enrichR (see Methods) scaled across all major cell classes for the top three differentially enriched (ordered by Combined Score, Methods) gene ontology terms for eight major cell classes. Terms colored red are the most enriched in the DA neuron class. **f**, Heatmap as in (**e**), for each of the 10 DA subtypes. Terms colored red are the most enriched in the *SOX6/AGTR1*+ population. **g**, Dot plot depicting the magnitude of disease-associated differential expression of selected genes in the “transcriptional from RNA polymerase II in response to stress” ontology term. **h**, Enrichment scores from GSEA (see Methods) for a p53-associated gene list^66^ (see Methods) for 10 DA subtypes. Color of lines represent the MASC odds ratio shown in Figure 2b.

A total of 26 genes have been identified within families whose mutation is linked to highly elevated PD risk^44^. We examined the overlap between these large effect-size variants and markers specifically expressed in the eight major cell classes (Fig. 3C). Consistent with our common variant analysis, DA neuron genes carried the only statistically significant (Bonferroni-corrected p-value < 0.05, Fisher’s exact test, Methods) burden of rare variant genes (Fig. 3C), suggesting that many of these variants act within DA neurons to influence neurodegeneration.

## Upregulation of p53-associated programs in the degenerating dopaminergic subtype

Finally, we used our profiling data to ask how the DA neurons alter their gene expression in response to PD. When compared to differential expression within the other major cell classes, DA neurons had the highest number of uniquely up- and down-regulated genes (Fig. 3D, Extended Data Fig. 8B, Methods). To gain insight into what pathways might be activated specifically in DA neurons, we performed Gene Ontology enrichment analyses on the significantly up-regulated genes across major cell types (Extended Data Fig. 8C, Methods and Supplementary Table 2). The most enriched ontologies associated with upregulated genes in DA neurons were the terms “regulation of neuron death” and “regulation of inclusion body assembly” (Fig. 3E). These ontologies were specifically enriched in the DA neurons (Fig. 3E), suggesting that the differential expression analysis was able to identify cell-specific regulatory changes expected in the context of neurodegeneration. The genes that drove this association included known regulators of cell death in PD and other diseases such as *DDIT4*^45,46^, *PPARGC1A*^47^, and *EGLN3*^48^, and others with as-yet unclear functions in the context of PD such as *RRAS2, TANK*, and *EFNB2* (Extended Data Fig. 8D).

To identify processes specific to our identified susceptible population, we performed gene ontology enrichments on DA subtype-specific differentially expressed genes (Extended Data Fig. 8E, Supplementary Table 3). The most significantly-enriched ontology term in the *SOX6/AGTR1* population was “transcriptional response by RNA polymerase II to stress” (Fig. 3F). The regulation of RNA polymerase II induced transcripts has been implicated in the response to excessive unfolded proteins^49^, a pathological hallmark of PD. We find *TP53* contributes to the enrichment of this ontology term and is also differentially expressed in the *AGTR1* subpopulation (Fig. 3G, Extended Data Fig. 8F). The protein p53 has been linked to PD-associated degeneration in mouse models, and itself can be regulated by parkin, a protein genetically linked to familial PD^50–53^. A heritability enrichment analysis suggests that an extended list of p53 targets (Methods) are significantly enriched in genes linked to PD common variation, at levels comparable to those associated with genetic risk for prostate cancer^54^ (Extended Data Fig. 8G). Finally, gene set enrichment analysis^55^ (GSEA) identified p53-regulated genes as most strongly and significantly (Bonferroni-corrected p-value < 0.01, Methods) enriched in the degenerating *SOX6/AGTR1*+ subpopulation (Fig. 3H), thereby nominating a potential pathway of neurodegeneration *in vivo*.

## Conclusions

In this study, we developed and applied a protocol for high-throughput expression profiling of individual human midbrain DA neurons. Our integrative analysis identified ten populations of SNpc neurons, including one that was primate-specific, adding to the short list of recently identified primate-specific brain cell types^23^. The characterization of differentially vulnerable DA populations provides several new scientific opportunities in the study of PD. First, the selective enrichment of expression of genes associated with PD genetic variants in the highly vulnerable *AGTR1* cells suggests that the pathways nominated by common variant studies may act cell-autonomously in these cells to influence neurodegeneration. Additional profiling of other cell types that are also vulnerable to PD-associated degeneration^56^ could clarify whether loss outside the SNpc is also cell-autonomous, or secondary to DA neuronal loss. Furthermore, our work should help refine *in vitro* DA neuron differentiation protocols, which could be used to screen for neuronal susceptibility across many different genetic backgrounds^57^ and with candidate therapeutic molecules. Xenotransplantation systems could also be useful to identify molecular regulators of neuronal vulnerability^58^, or to improve the performance of cellular replacement therapies^59^. Ultimately, such refined models and screening systems, ones that more closely match the molecular events occurring in affected patients, could prove to be useful tools in the understanding of a currently incurable disease.

## Acknowledgments

We would like to thank Djordje Gveric (Multiple Sclerosis and Parkinson’s Tissue Bank), Randall Woltjer (Oregon Health and Sciences University), and Rashed Nagra (Human Brain and Spinal Fluid Resource Center) for their contributions of postmortem tissue. We also thank Nicole Shultz and David Fitzpatrick (Max Planck Florida Institute for Neuroscience) for their donation of tree shrew brains. This work was supported by an NIH/NIA fellowship award to T.K. (1F30AG069446-261 01), an NIH New Innovator Award to E.Z.M. (DP2AG058488), an NIH/NIMH BRAIN Grant (1U19MH114821) to EZM, the Chan Zuckerberg Initiative (2017-175259) to E.Z.M., and the Fidelity Biosciences Research Initiative.

## Author Contributions

T.K. performed all analyses, with help from V.G. and A.A. A.A. developed the *NR4A2*-based nuclei enrichment protocol and generated the snRNA-seq data, with help from C.V. N.N. and C.V. generated the mouse midbrain dataset. S.B. and T.K. performed and analyzed the smFISH experiments. T.K. and E.Z.M. wrote the paper, with contributions from all authors.

## Methods

### Animal housing of *Mus musculus*

Animals were group housed with a 12-hour light-dark schedule and allowed to acclimate to their housing environment for two weeks post arrival. All procedures involving animals at MIT were conducted in accordance with the US National Institutes of Health Guide for the Care and Use of Laboratory Animals under protocol number 1115-111-18 and approved by the Massachusetts Institute of Technology Committee on Animal Care. All procedures involving animals at the Broad Institute were conducted in accordance with the US National Institutes of Health Guide for the Care and Use of Laboratory Animals under protocol number 0120-09-16.

### Brain preparation prior to 10x nuclei sequencing for *Mus musculus*

At 60 days of age, C57BL/6J mice were anesthetized by administration of isoflurane in a gas chamber flowing 3% isoflurane for 1 minute. Anesthesia was confirmed by checking for a negative tail pinch response. Animals were moved to a dissection tray and anesthesia was prolonged via a nose cone flowing 3% isoflurane for the duration of the procedure. Transcardial perfusions were performed with ice cold pH 7.4 HEPES buffer containing 110 mM NaCl, 10 mM HEPES, 25 mM glucose, 75 mM sucrose, 7.5 mM MgCl_2_, and 2.5 mM KCl to remove blood from brain and other organs sampled. The brain was removed immediately and frozen for 3 minutes in liquid nitrogen vapor and moved to −80°C for long term storage.

### Cryosectioning and brain preparation of postmortem frozen human brain

Post mortem human midbrain and dorsal striatum (caudate nucleus) tissue were flash frozen at −80 °C and cryosectioned at −15 to −20 °C into 60-micron sections. For midbrain preparation, the pigmented regions of the human midbrain were micro-dissected. Approximately 5-10 60-micron sections were made per frozen tissue block. Following microdissection, samples were placed on dry ice until nuclei isolation.

### Generation of single-nuclei suspensions from postmortem frozen human brain

To each cryosectioned sample, 1 mL of Extraction Buffer (ExB) was added into a 1.5-mL Eppendorf tube. Samples were briefly triturated before being placed in a six-well plate. Samples were then triturated 20 times with the ExB, every 2 minutes, until no large chunks of tissue were observed in each well. After the last trituration, samples were diluted with 45-50mL of wash buffer in a 50-mL Falcon tube, and then split into four 13-15 mL solutions in 50mL Falcon tubes. The diluted samples were then spun at 500g for 10 minutes at 4°C (pre-cooled) in a swing bucket benchtop centrifuge.

After centrifugation, a visible nuclei pellet was observed. Samples were then removed very gently from the centrifuge, and placed in an ice bucket. The supernatant was aspirated until there was barely any liquid observed on top of the pellet (50-100μL of liquid left). To aspirate without disturbing the pellet, a serological pipette was first used till about 1mL was remaining, followed by serial aspiration with a P2000 and P200 pipette.

The pellets were then resuspended in 250μL of wash buffer and mixed thoroughly by trituration. This 250μL solution was then pooled with four other pellets from the same original midbrain and resuspended into approximately 1mL of WB in an Eppendorf 1.5-mL tube.

### Immunolabeling and blocking of human and macaque nuclei

Approximately 100μL of 10% bovine serum albumin (BSA) in WB (final concentration of 1%) was added to the concentrated nuclei. Following the addition of a blocking solution, the anti-NR4A2-A647 (Santa Cruz, sc-376984 AF647) was then added at a concentration of 1:350. One caudate library was stained and sorted with NeuN (see metadata at Broad SCP link in Data Availability), which was performed by adding an anti-NeuN-PE (EMD MilliPore Corp., FCMAB317PE) antibody at a concentration of 1:1500. All samples were then wrapped with aluminum foil, and incubated on a rotator (lowest rotation speed) at 4°C for 0.5-1 hour. Following incubation, Eppendorf tubes were spun at 150g, for 10 minutes in a swing bucket benchtop ultracentrifuge (pre-cooled to 4°C). A visible pellet was observed after spinning. The supernatant was then gently aspirated, leaving almost no liquid. WB was then added to bring the total volume to approximately 1mL of sample. The samples were then stained with DAPI (ThermoFisher, #62248) at 1:1000 dilution. Samples were then filtered using a 70-micron cell filter before flow sorting.

### Fluorescence-activated nuclei sorting (FANS) for enrichment of dopaminergic nuclei

Flow sorting parameters for DAPI gating are described in Martin *et al* (https://www.protocols.io/view/frozen-tissue-nuclei-extraction-for-10xv3-snseq-bi62khge). For the NR4A2 positive selection on a flow sorter, a DAPI vs 647 gating was established by selecting the 2.5%-4% highest fluorescent *NR4A2* nuclei.

### Generation of single-nuclei suspensions from *Mus musculus, Rattus Norvegicus, Macaca mulatta*, and *Tupaia belangeri*

Frozen brains were carefully bisected sagitally with a cold (−20°C) razor blade. Each hemisphere was securely mounted by the lateral-most cortex onto cryostat chucks with OCT embedding compound such that the mid-brain was left exposed and thermally unperturbed. Dissection of the dopaminergic spans of the mid-brain including the substantia nigra pars compacta and reticulata as well as the ventral tegmental area was performed by hand in the cryostat using an ophthalmic microscalpel (Feather safety Razor #P-715) precooled to −20°C and donning 4x surgical loupes. Each excised tissue dissectate was placed into a pre-cooled 0.25 ml PCR tube using pre-cooled forceps and stored at −80°C. In order to assess dissection accuracy, 10um coronal sections were taken post dissection and imaged following Nissl staining.

Nuclei were isolated from frozen brain samples using a previously published^60^ protocol (https://www.protocols.io/view/frozen-tissue-nuclei-extraction-for-10xv3-snseq-bi62khge). See protocols.io link for all buffers and solution concentrations. All steps were performed on ice or cold blocks and all tubes, tips, and plates were precooled for >20 minutes prior to starting isolation. Briefly, 60 um sections of midbrain (~50 mg) were placed into a single well of a 12-well plate and 2 ml of ExB was added to each well. Mechanical dissociation was performed through trituration using a P1000 pipette, pipetting 1ml of solution slowly up and down with a 1ml Rainin (tip #30389212) without creating froth/bubbles a total of 20 times. Tissue was let rest in the buffer for 2 minutes and trituration was repeated. A total of 4-5 rounds of trituration and rest were performed (~10 minutes). The entire volume of the well was then passed twice through a 26-gauge needle into the same well. Following observation of complete tissue dissociation, ~2 ml of tissue solution was transferred into a precooled 50 ml falcon tube. The falcon tube was filled with Wash Buffer (WB) to make the total volume 30mls. The 30ml of tissue solution was then split across 2 different 50 ml falcon tubes (~15 ml of solution in each falcon tube). The tubes were then spun in a precooled swinging bucket centrifuge for 10 minutes, at 600 g at 4°C. Following spin, the majority of supernatant was discarded (~500μl remaining with pellet). Tissue solutions from 2 falcon tubes were then pooled into a single tube of ~ 1000μl of concentrated nuclear tissue solution. DAPI was then added to solution at manufacturer’s (ThermoFisher Scientific, #62248) recommended concentration (1:1000).

Flow sorting of isolated nuclei was performed according to the protocols.io link above. Briefly, 0.2ml PCR tube was coated with 5% BSA-DB solution. Solution was then removed and 20ul of FACS Capture Buffer was added as cushion for nuclei during sort. Nuclei were sorted into a chilled 96 well FACS plate (Sony M800 FACSorter). Sorting was done at a pressure of 6-7, with forward scatter gain of 1% on DAPI gate. The “purity” mode was used, and no spinning was performed after flow sorting nuclei into PCR tubes. Following sort, nuclei concentration was counted using a hemocytometer before loading into the 10X Genomics 3’ V3 Chip.

### Single-nucleus RNA-sequencing and library preparation

For all single-nuclei experiments, the 10X Genomics 3’ (V3) kit was used according to the manufacturer’s protocol recommendations. Library preparation was performed according to the manufacturer recommendations. Libraries were pooled and sequenced on either an Illumina NextSeq 500 or Illumina NovaSeq6000.

### Hybridization Chain reaction (HCR) on human and mouse tissue sections

Post mortem human and mouse midbrain tissues flash frozen in −80°C were cryosectioned at −15 to −20°C to make 12-micron sections on SuperFrost Plus slides. The slides are frozen at −80° until staining. After removal from −80°, the slides were allowed to warm up to room temperature (RT) before being placed in 4% PFA for 15 minutes at RT. Slides were then washed three times with 70% ethanol for 5 minutes before incubation at 70% ethanol for 2 hours at RT. After incubation, slides were then incubated at 37°C in the Probe Hybridization buffer (Molecular Instruments) for 10 minutes in a humidified chamber. The probe solution was then freshly prepared by adding 0.4 pmol of each probe set (Molecular Instruments) per 100 μL of Probe Hybridization buffer and vortexed. The Probe Hybridization buffer was then replaced by the probe solution. Slides were then incubated overnight at 37°C in a humidified chamber. After 18-24 hours, sections were sequentially washed in the following solutions: (1) 75% probe wash buffer and 25% 5x SSCT (SSC + 10% Tween-20), (2) 50% probe wash buffer and 50% 5x SSCT, (3) 25% probe wash buffer and 75% 5X SSCT, and (4) 100% 5x SSCT. The slides were then washed for 5 minutes at room temperature in 5x SSCT. Slides were then allowed to preamplify in the Probe Amplification buffer (Molecular Instruments) for 30+ minutes at RT. The hairpins were then freshly prepared by adding 1uL of hairpin per 100uL of Amplification Buffer (Molecular Instruments), and snap-frozen using a thermocycler with the following settings: 95° for 90 seconds, cool to room temperature (20°C) at a rate of 3 degrees per minute. Snap-frozen hairpins were added to the desired volume of the amplification buffer.

Slides were allowed to incubate overnight at RT in a humidified chamber. After overnight incubation, the slides are washed twice for 30 minutes at room temperature with 5x SSCT twice. An appropriate amount of Fluoromount Gold with NucBlue is added to the slides and then are coverslipped and sealed with clear nail polish. Slides were stored at 4°C till imaging.

| Probe Name | Accession Number |
| --- | --- |
| *TH* | NM_000360.4 |
| *CALB1* | NM_001366795 |
| *TMEM200A* | NM_001258276.1 |
| *SOX6* | NM_001145811.2 |
| *AGTR1* | NM_000685 |
| *GEM* | NM_005261.4 |
| *Th* | NM_009377.2 |
| *Calb1* | NM_009788.4 |
| *Gem* | NM_010276.4 |

### Imaging and analysis of single-molecule FISH experiments

Imaging was performed with a Nikon Eclipse Ti microscope and a Yokogawa CSU-W1 confocal scanner unit with an Andor Zyla 4.2 Plus camera. Images were acquired using a Nikon Apo 40x/1.15 WI objective.

For *in situ* validation of dopamine subtype vulnerability, slides were viewed in their entirety by scanning the tissue with the 40x water objective (Nikon). An area was considered to contain a single-positive neuron if the following criteria were met: neuron is at least 70% in frame, there is DAPI signal within the neuron, there is signal in the 488nm channel that does not overlap with other channels, and distinct puncta are visible. An area was considered to contain a doublepositive neuron if the previous criteria were met and the neuron contains at least 3-5 distinct, non-overlapping puncta in the 561nm channel. An area is considered to contain a triple-positive neuron if all the previous criteria is met and the neuron contains at least 3-5 distinct, nonoverlapped puncta in the 647nm channel.

A total of 10 control and 10 Parkinsonian midbrains were imaged and quantified for the following sets of markers: *TH/AGTR1/SOX6* and *TH/CALB1/TMEM200A*. All manual quantification of subpopulations was performed blinded to case versus control status for both probe sets. To generate the p-values in Fig. 2H, a Wilcoxon rank-sum test was performed on the fractional abundance of double- and triple-positive cells.

### Generation of slides, imaging, and image analysis for stereotactic localization of dopaminergic subtypes

To generate a stereotactic map of the select transcriptionally-defined midbrain dopaminergic neurons, we obtained one fully intact postmortem brainstem/midbrain from one neuropathologically/neurotypically-normal 32-year old male with the entire substantia nigra pars compacta present bilaterally. One hemisphere of the midbrain was used to excise the pars compacta in its entirety, creating a block (approximately 2cm by 3cm by 0.43cm) which was then mounted to a cryostat chuck with OCT embedding compound and frozen at −20 °C. The entire block of tissue was then serially sectioned at 12 micrometers, each section being placed on an individual SuperFrost Plus (VWR) glass slide. The anterior to posterior estimate was carefully recorded, and slides were thawed in groups of six, equally spaced. Hybridized Chain Reaction (HCR) was performed as above on two select groups of markers: *TH, CALB1*, and *TMEM200A* and *TH, SOX6*, and *AGTR1* (see above for accession numbers).

Imaging was performed with a Nikon Eclipse Ti microscope and a Yokogawa CSU-W1 confocal scanner unit with an Andor Zyla 4.2 Plus camera. Images were acquired using a 20x air objective (Nikon). Images were acquired with 80% laser power and 300ms exposure. The number of fields to be imaged in the x and y axis was determined by manually testing various amounts until the entire visible substantia nigra was contained within the area to be imaged.

Each image was converted into a composite image and, using the criteria defined above for *in situ* validation of proportional alterations, were sorted depending on whether or not they contained neurons using FIJI. A binary mask was applied to the images based on thresholding intensity, area, and circularity (representative in Extended Data Figure 6B). The 3D Objects Counter function in FIJI was applied to the binary masks for each image and the results were saved and compiled into a single Excel sheet. Each detected mask was then multiplied by the other two channels and cells were manually annotated as single-positive (*TH+/CALB1- /TMEM200A-* or *TH+/SOX6-/AGTR1-*) or triple-positive (*TH+/CALB1+/TMEM200A*+ or *TH+/SOX6+/AGTR1*+). A minimum volume value of 5000 from the 3D Objects Counter analysis in order to remove auto fluorescent puncta and sections of DA neurons that were partially cut.

To generate the images in Figure 2E and Extended Data Figure 6A, the masks and their associated label (single- or triple-positive neurons) were aggregated across all tiles for each of the six slides imaged and colored using the ggplot function in R. To generate the interpolated plots in Figure 2F, the six plots were registered and merged by scaling and centering the x- and y-values. Then, the values were interpolated in 3D using the interp function in the akima package in R: https://cran.r-project.org/web/packages/akima/index.html. Interpolated values were displayed using the persp3D function in the plot3D package in R: https://cran.r-project.org/web/packages/plot3D/index.html.

### Pre-processing of sequencing reads

Sequencing reads from human midbrain experiments were demultiplexed and aligned to the hg19 reference using DropSeqTools (https://github.com/broadinstitute/Drop-seq) with the default settings, including exonic and intronic data as UMIs. Gene expression matrices from the human midbrain experiments were generated using DropSeqTools. Reads from all non-human species were aligned using DropSeqTools against the following genomes: *rattus norvegicus* (Rnor_6.0), *mus musculus* (mm10), *macaca mulatta* (Mmul_10), and *tupaia belangeri* (tupBel1). Digital count matrices were subsequently generated from DropSeqTools. Sequencing reads from human caudate nucleus experiments were demultiplexed and aligned to the hg38 reference using CellRanger v3 with default settings. Counts were generated using the CellRanger “count” function.

### Cell-type clustering and annotation

Cell types were defined using a two-step process. First, individuals were clustered independently to extract major cell types using a modified version of the Seurat (v2)^61^ workflow. Once major cell types were extracted, subtypes were defined using a joint integration with either Harmony^62^ for non-neuronal cell types or LIGER’s projection method for neuronal cell types.

We used Harmony to integrate non-neuronal cell types across individuals into a shared space by removing batch effects while preserving the biological variation. The following analysis steps were performed for each non-neuronal cell type, separately. First, we searched for highly variable genes in each of 18 individual-level datasets using a variance stabilizing transformation method from the Seurat package in R. Genes identified as highly variable in at least 4 individual-level datasets were selected for principal component analysis (PCA). Next, individual-level datasets were combined after standardizing the expression of highly variable genes to have a mean of zero and variance of one. We performed PCA analysis with PCs weighted by the variance that they explained using Seurat. To integrate datasets, a grid search was performed on the number of PCs and the harmony parameter sets to find a solution with optimal mixing of the cells from different subjects, while maximizing the separation of different cell states as judged by the expression patterns of marker genes. The integrated space was used to construct the nearest neighbor graph. Clustering was performed using the SLM algorithm with a resolution of 0.8.

For neuronal cell types, the LIGER projection method was used in order to minimize the influence of the diseased cell transcriptome on the reduced space. Briefly, neuronal nuclei from the eight neurologically normal individuals were clustered and annotated using the standard LIGER workflow. Variable genes were selected followed by integrative non-negative matrix factorization and quantile normalization. Next, profiles from individuals with PD/LBD were integrated by projecting nuclei onto the iNMF dimensions generated from the control individuals. Clusters were defined by the SLM algorithm at a resolution of 0.6 and marker genes were generated using a Wilcoxon rank-sum test comparing nuclei from one cluster against the rest of the nuclei. The alignment score calculations in LIGER were performed using the default package settings.

### Identification of specific midbrain dopaminergic neuron transcription factors

To nominate potential nuclear transcription factors (TFs) for flow-based enrichment (Extended Data Figure 1A), we used a recently-published comprehensive survey of the mouse brain^20^. We downloaded midbrain data and performed differential expression between DA neurons and all other cell types using a Wilcoxon rank-sum test from the Presto package (https://github.com/immunogenomics/presto). We intersected our list with a list of mouse TFs (http://genome.gsc.riken.jp/TFdb/cdimage/htdocs/tf_list.html) and determined the AUC (area under curve) using the default statistics from the Presto package.

### Integration of dopaminergic neurons across species

In order to jointly define dopaminergic neurons across species, we used LIGER’s projection method. Briefly, we subsetted DA neurons from all species based on the expression of *TH*. We first integrated one mouse and one human DA neuron dataset using LIGER with the following parameters: number of variable genes = 1,080, k = 10 (number of latent factors), lambda = 5 (strength of integration), knn_k (number of nearest neighbors) = 45, SLM resolution (number of clusters) = 1.5. We then projected on rat, tree shrew, and macaque DA neurons using the online iNMF branch of LIGER, with the setting “projection = TRUE”. After quantile normalization, datasets were jointly clustered using the SLM algorithm in LIGER with a resolution of 1.5. The joint low-dimension embedding was visualized with t-Distributed Stochastic Neighbor Embeddings (t-SNE).

For two-way integrations (mouse versus human and macaque versus human), we used the standard LIGER pipeline, selecting between 1000-2000 highly variable genes between human and mouse and human and macaque. We used the following parameters for both integrations: k = 10, lambda = 5, knn_k = 20, SLM resolution = 0.6.

### SCENIC analysis

To identify differentially regulated regulons associated with specific dopaminergic subtypes, we employed Single-Cell rEgulatory Network Inference and Clustering (SCENIC) with user recommended settings from the SCENIC vignette (http://htmlpreview.github.io/?https://github.com/aertslab/SCENIC/blob/master/inst/doc/SCENIC_Running.html). Briefly, we averaged the log-normalized gene values of single-nucleus profiles from our dataset of non-Parkinson’s control individuals per dopaminergic subtype. We filtered on genes that had at least one count per bulk profile across all 10 subtypes. We ran a correlation analysis using GENIE3 and then ran SCENIC to determine transcription factor modules within these correlations. Using AUCell, we scored cells based on regulon activity and plotted these scaled regulon scores on a per dopaminergic subtype basis. To determine statistically significant differences in regulon activity, we ran a Wilcoxon rank-sum test (using the presto package: https://github.com/immunogenomics/presto) on the regulon scores between major dopaminergic subtypes and ranked cells based in their area under the curve and an Benjamini-Hochberg adjusted p-value < 0.05.

### Differential abundance assessment of cell types in association with PD and other covariates

To identify differentially abundant cell subpopulations in association with PD, we employed MASC, or Mixed-effects Association of Single cells^32^. MASC is a generalized mixed-effect model with a binomial distribution which ascertains whether a certain covariate significantly influences the membership of a given nuclei. We included the following fixed covariates into the generalized mixed-effect model: *sex*, *status* (control or disease), and *NR4A2* (whether the nuclei were captured in a *NR4A2*-positive or *NR4A2*-negative library). Finally, the library from which the nuclei was sampled was included as a random effect to account for the intra-library correlation of cell numbers. Cell subpopulations were considered significant at FDR-adjusted p-value < 0.05 and an absolute odds ratio greater than 0.

### Disease enrichment embedding score

To project information about relative abundance onto the reduced dimension space for the DA subtypes, we developed an enrichment score (Fig. 2C). Briefly, we binned the two-dimensional embedding for both disease and control individuals. For each bin, we determined the total number of cells per individual within that bin. We then averaged those values across individuals per bin for both case and control. We then scaled these scores across all bins to provide a normalized estimate of the relative abundance of certain DA subtypes across the UMAP space.

### Differential expression and GO term enrichment

To identify differentially expressed genes across all major cell types, we employed MAST (Model-based Analysis of Single-cell Transcriptomes)^63^. MAST is a mixed-effect hurdle model that models droplet-based single-nucleus/cell expression data as a mixture of a binomial and normal distribution (for the log-normalized non-zero expression values) while systematically accounting for pre-defined covariates. We included the following fixed-effect covariates in our model: *log*(*number of UMIs*), *sex, percent.mito* (percent of reads that map to mitochondrial genes per nucleus) and *status* (control or disease) to test the effect of the disease on each cell type. We additionally included the library from which a nucleus was sampled from as a random effect covariate in the model to account for intra-library correlation of expression data. To evaluate the effect of disease on expression, we performed differential expression analysis across all major cell types by evaluating the significance of the fixed effect of disease status using a Wald test. For all non-dopaminergic cell types, we used only the cells from the NR4A2-negative libraries. For dopaminergic neurons, we used the *NR4A2*-positive gated libraries. Genes were defined as significantly differentially expressed at a Benjamini-Hochberg-adjusted p-value < 0.01. For all expression tests, we used the discrete (the binomial link part of the hurdle model) coefficient of MAST to determine the coefficient estimate of the effect of the disease on the expression.

To identify differentially expressed genes between clusters, we down-sampled the control datasets 100,000 nuclei and performed MAST comparing between cell types and subtypes while accounting for batch variation. We included all the same covariates as above (except for disease status) and included additional covariates of cell type identity and a covariate for whether the cell derived from a *NR4A2*-positive or *NR4A2*-negative library. To create the marker gene sets, we selected all genes with a MAST discrete coefficient (the binomial link part of the hurdle model) greater than 0 and a Benjamini-Hochberg-adjusted p-value of less than 0.01.

To identify differentially enriched gene ontology terms, we used enrichR^64^ and tested all GO categories within the GO Molecular Functions (2018) and Elsevier Pathway Collections. We used genes that were significantly differentially expressed per major cell type and subtype and identified significant ontologies at a Benjamini-Hochberg-corrected significance level of 0.1. Specificity was determined on a per ontology level with the scaled Combined Score.

### Heritability enrichment analysis

To determine which cell types and subtypes are enriched for heritable risk of traits, we used the package MAGMA^40^ (https://ctg.cncr.nl/software/magma). For Alzheimer’s disease (AD) and Parkinson’s disease, we downloaded publicly available summary statistics from the most recently available study^41,43^. We then performed a SNP to GENE calculation with either the MAGMA tool or the equivalent webserver (FUMA: https://fuma.ctglab.nl/) using default parameter settings. For all major cell types/subtypes, we took the top 3500 marker genes ranked by the Z-score determined from MAST (see above section) for each cell type and subtype as a defined gene set. As an alternate approach for marker selection per cell type, we defined as the top 10% of genes ranked by their area under the curve (AUC) per major cell type, an approach used by previous efforts to nominate cell types using heritability enrichment analyses ^65^. We ran MAGMA on the set annotation setting to test the significance of whether the marker gene set of that cell type was enriched for heritable risk of the trait. The resulting p-values were corrected for multiple hypothesis testing across major cell types and subtypes with the Bonferroni method (8 tests for major cell types and 68 tests for subtypes).

For testing significance of p53 targets for enrichment of genetic risk, we used a published list of p53 targets^66^ and ran MAGMA on the default “set annotation” settings for AD, PD, schizophrenia (SCZ), and prostate cancer (PrCan). Schizophrenia^67^ and prostate cancer^54^ summary statistics can be downloaded at: https://www.med.unc.edu/pgc/download-results/ and https://www.ebi.ac.uk/gwas/publications/29892016.

### Parkinson’s disease familial variants enrichment analysis

To test whether there exists any enrichment of genes that have been previously nominated as containing variants that cause familial Parkinson’s disease, we gathered a list of known familial PD genes from a recent whole-exome sequencing study^44^. We ran a Fisher’s exact test using the geneOverlap package between genes that were considered specifically expressed in a major cell type, as determined by the top 10% of genes ordered by the AUC metric with the Wilcoxon rank-sum test, and these 26 genes. The resulting p-values were then corrected for multiple hypothesis testing using the Bonferroni method (8 tests for major cell types).

### Gene Set Enrichment (GSEA) analysis of p53 gene sets on dopamine subtypes

GSEA was performed using the fGSEA package^68^ using the following settings: minSize = 1, maxSize = 450, and nperm = 10000. The p53 gene set was defined from a previous survey of p53 targets^66^. Enrichment scores were calculated from fGSEA and all p-values were corrected for multiple hypothesis testing with the Bonferroni method (10 tests for dopamine subtypes).

### Robustness testing of specificity of differentially expressed genes in dopaminergic subtypes

In order to test the robustness of the specificity of differentially expressed genes (Extended Data Fig. 8F) between subtypes, we sampled all non-*SOX6+/AGTR1*+ cell types and aggregated them together. We then ran MAST (see above) on a combined group of DA neurons and compared the coefficient and p-values with the original and downsampled versions of the *SOX6+/AGTR1*+ differential expression analysis. All p-values were corrected for multiple hypothesis testing using the Benjamini-Hochberg adjustment.

### Statistics and Reproducibility

All statistical tests are performed using a two-sided test, unless noted above. All statistical tests were corrected for multiple hypothesis testing using either a Benjamini-Hochberg adjustment or Bonferroni correction (specific choices are listed in the above methods).

### Data Availability

All processed data, along with UMAP coordinates for major integrations, are available at: https://singlecell.broadinstitute.org/single_cell/study/SCP1402. Processed digital gene expression (DGE) matrices discussed in this publication have been deposited in NCBI’s Gene Expression Omnibus and are accessible through GEO Series accession number GSE178265 (https://www.ncbi.nlm.nih.gov/geo/query/acc.cgi?acc=GSE178265). Raw sequencing data associated with all snRNA-seq experiments will be made available upon publication; a dbGAP deposition is pending. Raw count and proportion values for the *in situ* validation experiment can be found in Supplementary Table 1.

---

**Extended Data Figure 1:**
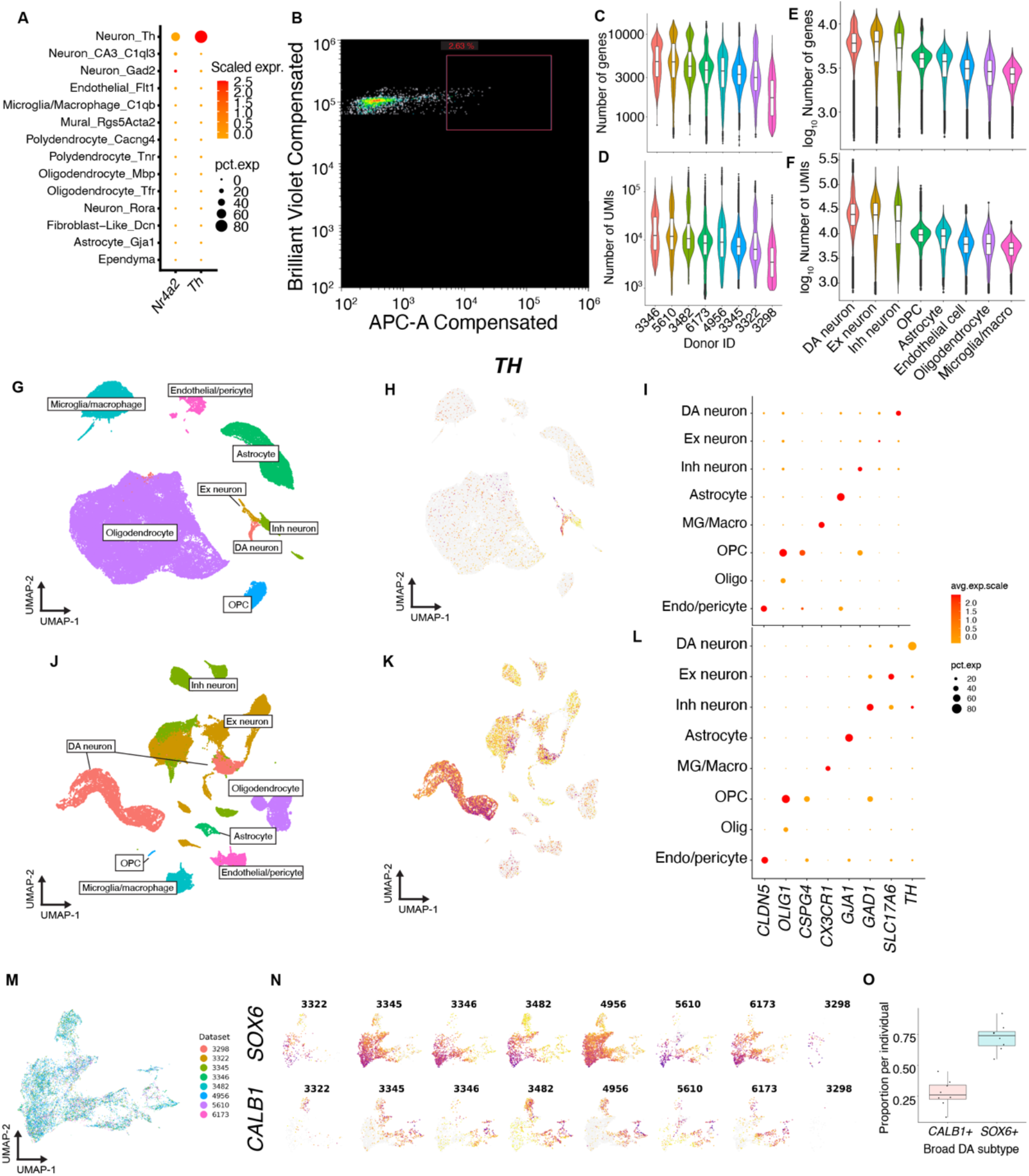
High-throughput snRNA-seq profiling of SNpc. **a,** Expression of tyrosine hydroxylase (*Th*) and *Nr4a2* in a published scRNA-seq study of mouse midbrain^20^ **b,** Representative FANS plot for enriching midbrain dopaminergic neurons. APC-A (x-axis) represents *NR4A2* antibody channel and Brilliant Violet (y-axis) represents DAPI channel. The NR4A2 gate was thresholded to select the top 2% of all nuclei (blue box). **c,** Number of genes per donor. **d,** Number of UMIs (unique molecular identifiers) per donor. **e,** log_10_ number of genes per cell type. **f,** log_10_ number of UMIs per cell type. **g,h,** UMAP representations of the NR4A2-gated nuclei, colored by cluster (**g**), and expression of *TH* (**h**). **i**, Dot plots of marker genes for the eight major cell classes in the NR4A2-gated nuclei. **j,k,l,** The same analyses shown in **g, h**, **i** but for the NR4A2+ gated nuclei. DA, dopamine; Ex, excitatory; Inh, inhibitory; Astro, astrocyte; MG, microglia/macrophages; OPC, oligodendrocyte precursor cell; Olig, oligodendrocyte; Endo, endothelial cells), **m,** UMAP representation of 15,684 DA profiles colored by individual. **n**, Top - expression of *SOX6* per individual. Bottom - expression of *CALB1* per individual. **o,** Box plot of proportion of *SOX6*+ and *CALB1*+ DA neurons per individual. Median is defined by the center line; interquartile ranges are shaded by broad subtype. Dots represent individual samples.

**Extended Data Figure 2:**
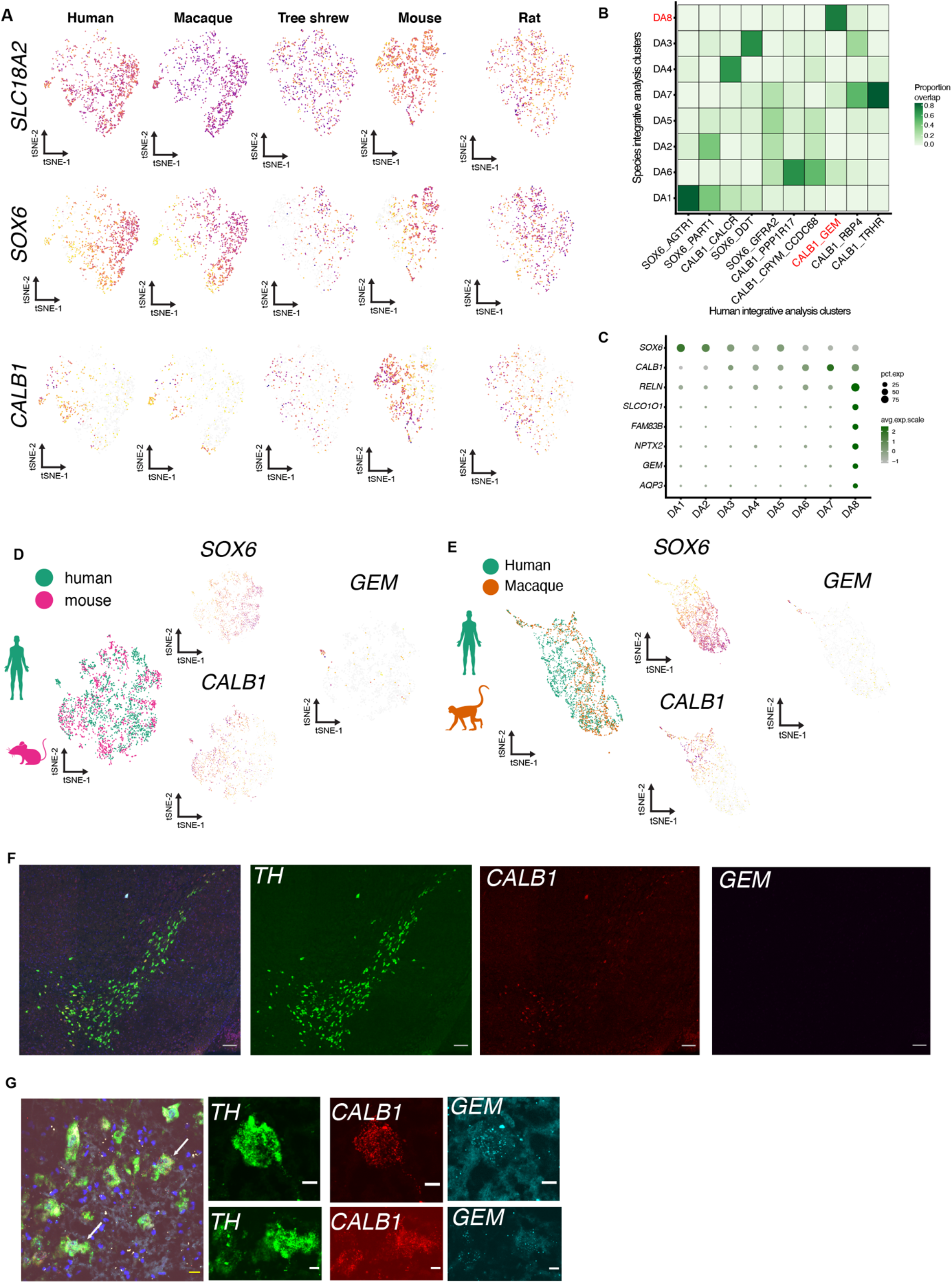
Cross-species analysis of DA subtypes. **a,** Expression of *SLC18A2* (DA neuron marker, top row), *SOX6* (middle row), and *CALB1* (bottom row) across all five species sampled. **b,** Confusion matrix showing overlap of human cells within clusters defined by the cross-species integrative analysis and the human-only analysis (**Fig. 1**). The clusters colored in red correspond to the primate-specific population (DA8) and cognate population in the human-only analysis (CALB1_GEM). **c,** Dot plot of additional marker genes for the primate-specific population, DA8. **d,** UMAP representation of two-way integrative analysis between mouse and human colored by species (left), and major cell markers plotted (top-center = *SOX6*, bottom-center = *CALB1*, and right = *GEM*). **e,** Two-way integrative analysis between macaque and human colored by species (left) and major cell markers plotted (top-center = *SOX6*, bottom-center = *CALB1*, and right = *GEM*). **f,** Tiled images of mouse midbrain for marker genes *Th, Calb1*, and *Gem* (grey scale bar = 100 μm). **g,** Additional images of *TH+/CALB1+/GEM*+ cells (white arrows) within human midbrain (yellow scale bar = 20 micrometers, white scale bar = 10 μm).

**Extended Data Figure 3:**
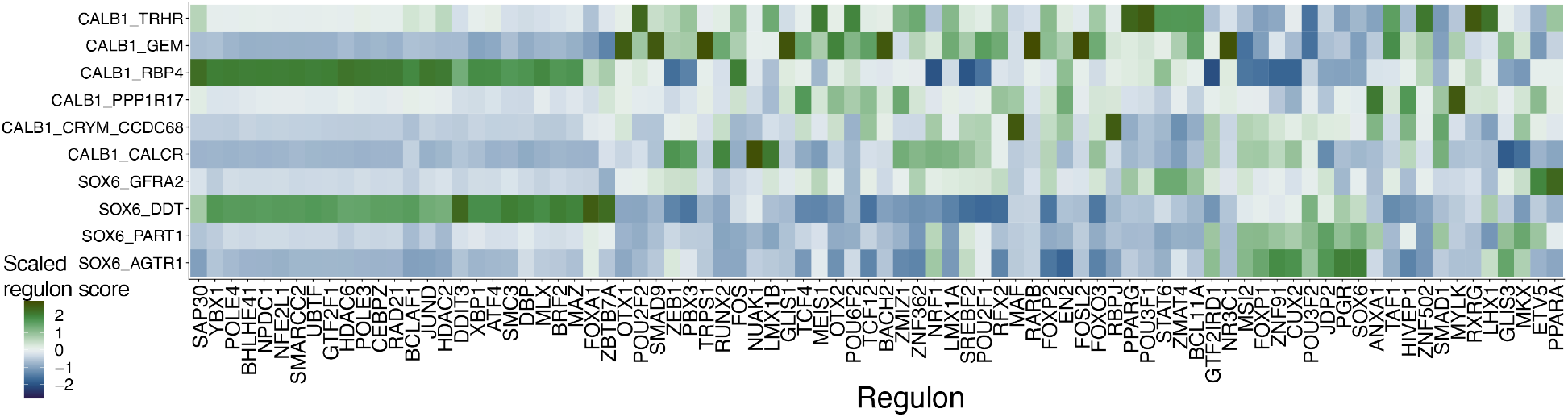
Differentially expressed regulons across all DA subtypes. Differentially expressed (AUC > 0.7, FDR-adjusted p-value < 0.05) regulons for all 10 DA subtypes. Scaled regulon score is the regulon activities as determined from SCENIC analysis (see Methods) averaged per subtype and scaled across all 10 subtypes.

**Extended Data Figure 4:**
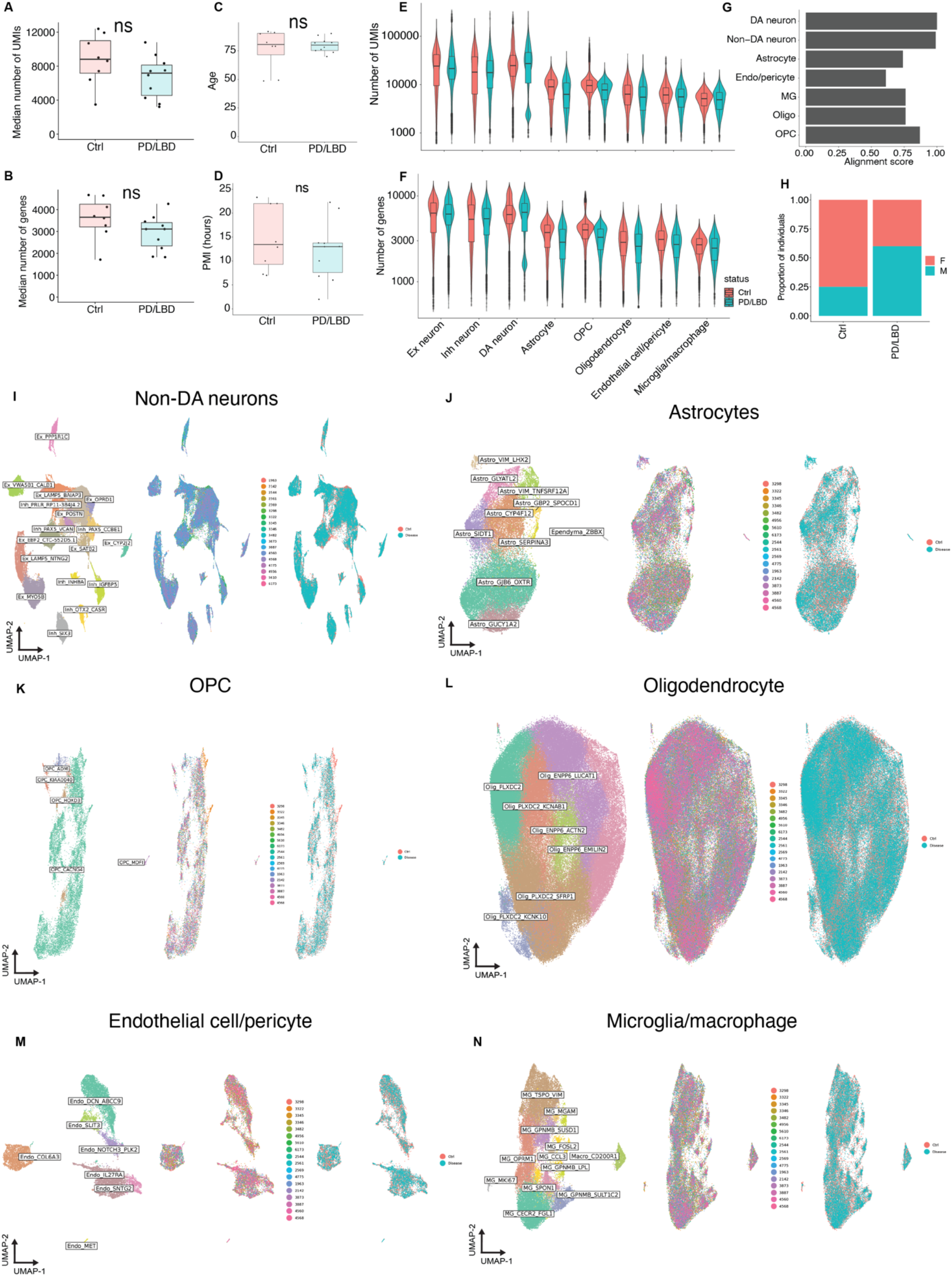
Case-control scRNA-seq integrative analysis of SNpc. **a,b,c,d,** Box plots showing, per individual, the median number of UMIs (**a**) and genes (**b**) per profile, age at death (**c**) and postmortem interval (**d**) stratified by case (PD/LBD) and control (Ctrl). ns, not significant Wilcoxon rank-sum test p > 0.05. For all boxplots, median is center line, interquartile ranges are shaded according to disease status, and dots represent values for each individual. **e, f** Violin plots of number of UMIs (**e**) and genes (**f**) per profile across eight major cell classes grouped by disease (PD/LBD, blue) and control (Ctrl, red). **g,** Alignment scores (see Methods) for each of the seven non-DA cell classes. **h,** Stacked bar plot of sex stratified by disease status. **i-n,** UMAP representations of Non-DA neurons (n = 91,479 nuclei) **(i)**, Astrocytes (n = 33,506) **(j)**, Oligodendrocyte precursor cells (OPCs) (n = 13,691) **(k)**, Oligodendrocytes (n = 178,815) **(l)**, Endothelial cells/pericytes (n = 14,903) **(m)**, and Microglia/macrophages (n = 33,041) **(n)** colored by cell type (left), individual (middle), and disease status (right).

**Extended Data Figure 5:**
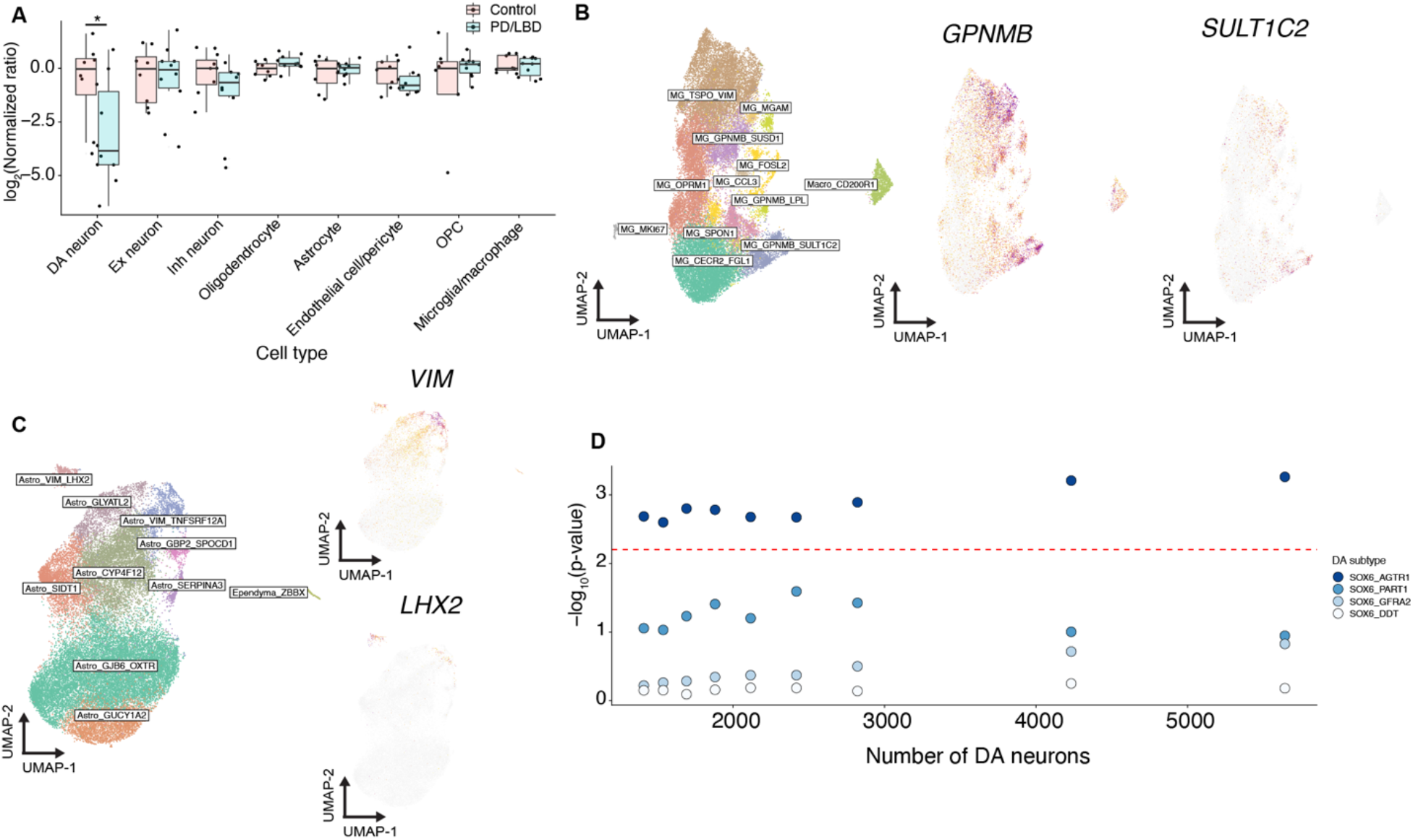
Analyses of cell type proportional changes in PD/LBD. **a,** Box plot of log_2_ ratio of cell types normalized to median of control ratios. Pink and blue dots denote control and PD/LBD individuals, respectively (* = p < 0.05, Wilcoxon rank-sum test). Median is defined by the center line, and interquartile ranges are shaded by control versus PD/LBD (see legend). **b,** Left = UMAP of 33,041 microglia/macrophage cells. Middle and right = expression of *GPNMB* and *SULT1C2* marking specific expression in region of annotated cluster. **c,** Left = UMAP of 33,506 astrocytes. Top and bottom right = expression of *VIM* and *LHX2* respectively marking specific expression in region of annotated cluster. **d,** Dot plot of downsampling analysis for *SOX6*+ DA subtypes. X-axis denotes size of dataset after downsampling and Y-axis is the MASC-computed −log_10_ p-values associated with disease status. Red dotted line is FDR-adjusted p-value = 0.05. Dots are colored by subtype.

**Extended Data Figure 6:**
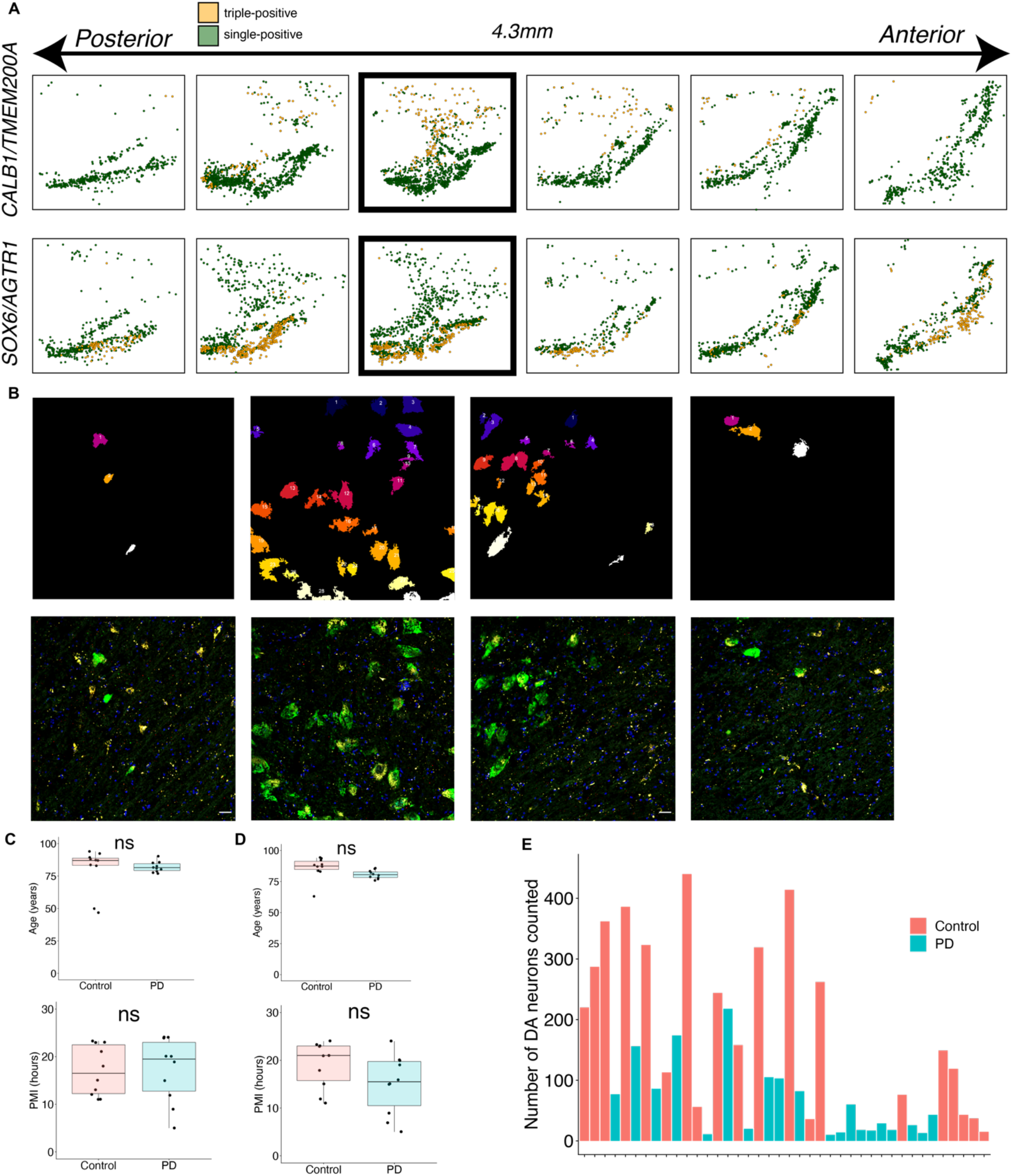
Imaging of highly vulnerable and resistant DA subtypes in the SNpc by smFISH. **a,** Stereotactic localization of *TH+/CALB1+/TMEM200A*+ neurons (top) and *TH+/SOX6+/AGTR1*+ neurons (bottom) across six coronal sections equally spaced along the 4.3mm of the rostral-caudal axis. The highlighted plates are also shown in Figure 2e. **b,** Top row, representative masks of segmented cells used to generate the quantification in (**a**). Bottom row, original images used to generate the masks above. **c**, **d**, Box plots, stratified by disease status, of age of death (top) and postmortem interval (bottom), for each subject in the *in situ* quantification of *TH+/CALB1+/TMEM200A*+ cells (**c**) and *TH+/SOX6+/AGTR1*+ cells (**d**). For both boxplots, medians are defined by center line, interquartile ranges are shaded by disease classification (control versus PD), and dots represent individual samples. **e,** Total number of DA neurons counted per slide; bars are colored by disease status.

**Extended Data Figure 7:**
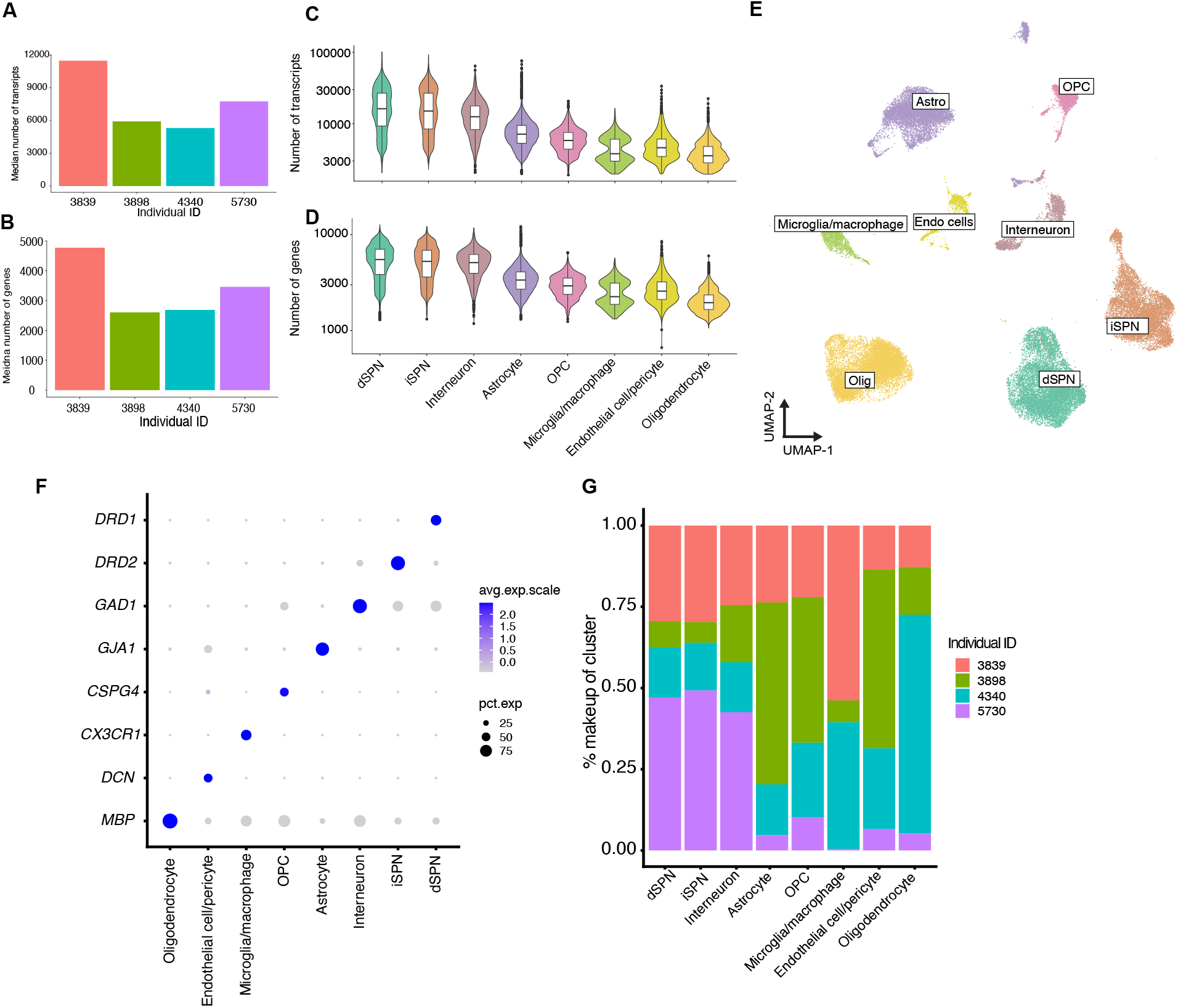
Analysis of human dorsal striatum by snRNA-seq. **a,b**, Median number of UMIs (**a**) and genes (**b**) per donor. **c,d** Violin plots showing the number of UMIs (**c**) and (**d**) across eight major cell classes defined by clustering. dSPN, direct spiny projection neurons; iSPN, indirect spiny projection neurons; OPC, oligodendrocyte precursor cells. **e,** UMAP representation of 46,872 single nuclei from the dorsal striatum colored by major cell class. Astro, astrocyte; Olig, oligodendrocyte; Endo cells, endothelial cell/pericyte. **f,** Dot Plot showing specific expression of selected marker genes across the eight major cell classes. **g,** Percent contribution of four tissue donors to the eight major cell classes.

**Extended Data Figure 8:**
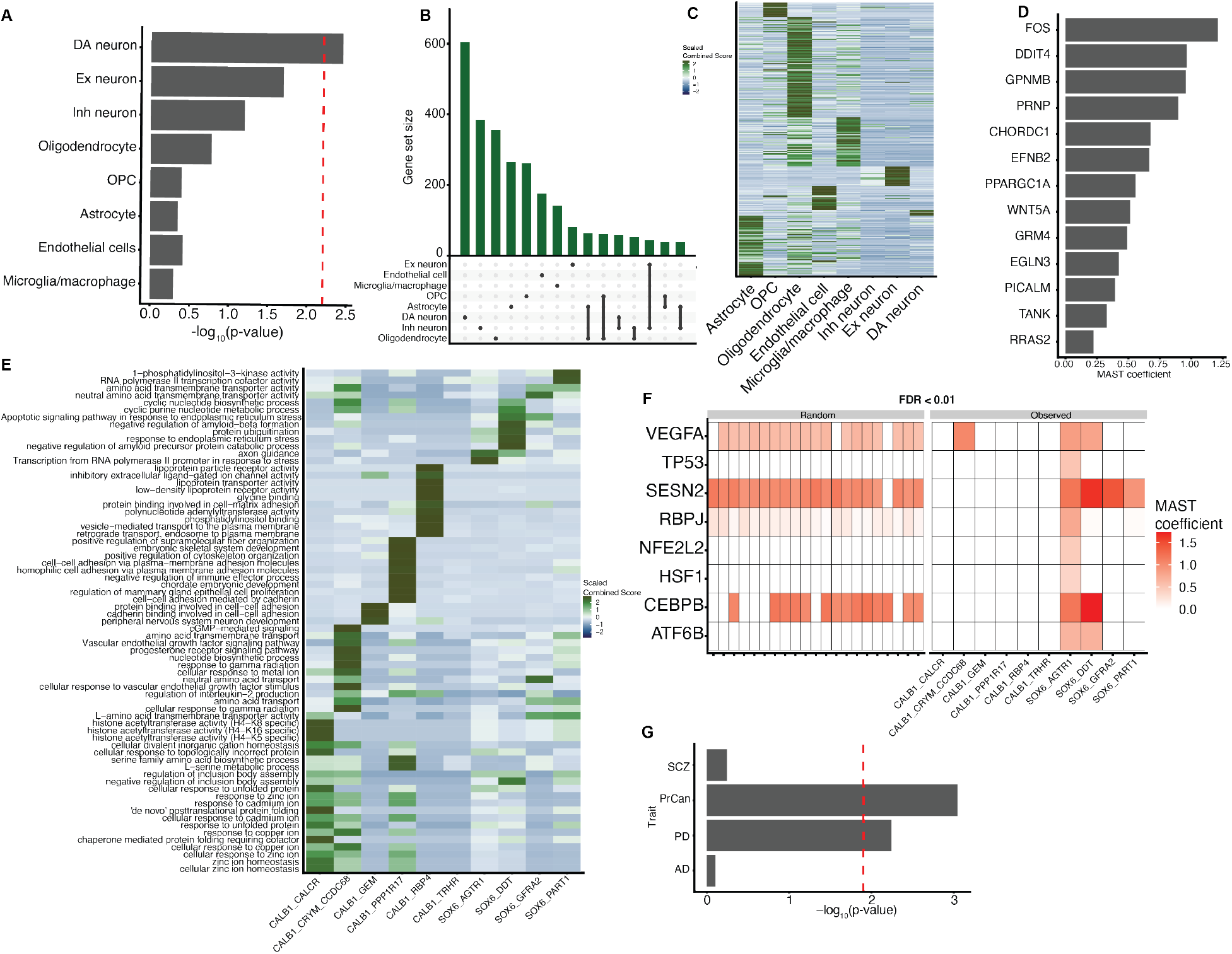
Robustness testing of genetic correlation and differential expression hypotheses. **a,** Bar plot of −log_10_-transformed p-values from MAGMA enrichment of PD across the eight SNpc cell classes using genes defined by Wilcoxon rank-sum AUC. Dotted red line indicates Bonferroni significance (p < 0.05/8). **b,** Upset plot of top 1000 down regulated genes across all eight major cell types in SNpc (Methods). **c,** Heatmap of scaled Combined Score (Methods) from all enriched Gene Ontology terms across eight SNpc cell classes in the PD/LBD cases. **d,** MAST coefficients (Methods) for genes that drive association of “regulation of neuron death” ontology term. **e,** Heatmap of scaled Combined Score (Methods) for all enriched Gene Ontology terms from all upregulated genes in the 10 DA subtypes. **f,** Heatmap showing robustness testing results (Methods) to confirm specificity of differential expression to *SOX6/AGTR1* DA subtype in PD. Colors of tiles are binarized MAST coefficient values (white if FDR-adjusted p-value > 0.01). Left = MAST binarized coefficients for genes driving “regulation of RNA polymerase II transcription” generated from randomly sampled *non-SOX6/AGTR1* DA subtypes. Right = MAST binarized coefficients from analysis on *SOX6/AGTR1* DA subtypes **g,** Bar plot of −log_10_-transformed p-values from MAGMA enrichment of extended p53 list^66^ over four traits (Methods, SCZ = schizophrenia, PrCan = prostate cancer, PD = Parkinson’s disease, AD = Alzheimer’s disease). Dotted red line indicate Bonferroni significance level (p < 0.05/4).

**Extended Data Table 1:** Basic demographics for 8 neuropathologically normal individuals used for snRNA-seq midbrain experiments

| Sample ID | Brain Bank | Age (yrs) | Sex | PMI (hrs) | Neuropathological evaluation | Cause of death |
| --- | --- | --- | --- | --- | --- | --- |
| 3322 | Sepulveda | 90 | F | 22 | No evidence of neurodegenerative disease | Colon cancer |
| 3298 | Sepulveda | 79 | M | 22 | No evidence of neurodegenerative changes by H&E or silver stain | Prostate cancer |
| 4956 | Sepulveda | 92 | F | 23.3 | Examination shows essentially normal brain | Uterine cancer |
| 3345 | Sepulveda | 82 | F | 12.9 | No gliosis or NFT present. Normal neuronal cellularity (neocortex/hippocampus) | Colon cancer |
| 3482 | Sepulveda | 79 | F | 14 | Examination reveals no pathology | Coronary artery disease |
| 3346 | Sepulveda | 91 | F | 10 | No gliosis or NFT present. Normal neuronal cellularity (neocortex/hippocampus) | Congestive Heart failure |
| 6173 | UMaryland | 49 | M | 13 | Unremarkable adult brain and dura mater | Not listed |
| 5610 | UMaryland | 50 | F | 11 | Essentially normal brain | Smoke inhalation |

**Extended Data Table 2:** Basic demographics for 10 PD/LBD individuals used for snRNA-seq midbrain experiments

| Sample ID | Brain Bank | Age (yrs) | Sex | PMI (hrs) | Neuropathological evaluation | Midbrain loss evaluation* | Cause of death |
| --- | --- | --- | --- | --- | --- | --- | --- |
| 4775 | Sepulveda | 75 | M | 13.8 | Parkinson's disease | "Moderate decrease" | Parkinson's disease |
| 3873 | Sepulveda | 76 | F | 6.8 | Parkinson's disease | "Prominent loss" | Parkinson's disease |
| 4560 | Sepulveda | 83 | F | 13 | Parkinson's disease | "moderate decrease" | Parkinson's disease |
| 3887 | Sepulveda | 82 | M | 13.7 | Parkinson's disease | "Moderate decrease" | Parkinson's disease |
| 4568 | Sepulveda | 78 | M | 22.3 | Parkinson's disease | "Prominent decrease in pigmented neuronal density" | Parkinson's disease |
| 2561 | OHSU | 70 | F | 21 | Lewy body dementia | 2 | Lewy body dementia |
| 2142 | OHSU | 90 | M | 2 | Parkinson's disease | 1 | Parkinson's disease |
| 2569 | OHSU | 74 | M | 13 | Lewy body dementia | 2 | Lewy body dementia |
| 2544 | OHSU | 89 | F | 10 | Lewy body dementia | 2 | Lewy body dementia |
| 1963 | OHSU | 82 | M | 6 | Parkinson's disease | 2 | Parkinson's disease |

\*Note: midbrain loss scores (if reported by brain bank) range from 0-3 (0 = none, 3 = severe)

**Extended Data Table 3:** Demographics and neuropathology for 10 control and 10 PD frozen midbrains used for in situ validation of selective vulnerability

| Subject ID | Age at death (years) | Sex | PMI (hours) | Status | Used in CALB1/TMEM200A set | Used in SOX6/AGTR1 set | Lewy body score (NP = not provided) |
| --- | --- | --- | --- | --- | --- | --- | --- |
| PDC107 | 87 | F | 15 | Ctrl | x | x | NP |
| C064 | 63 | F | 21 | Ctrl | x |  | NP |
| C087 | 94 | F | 24 | Ctrl | x |  | NP |
| C088 | 94 | M | 11 | Ctrl | x | x | NP |
| C043 | 87 | F | 12 | Ctrl | x | x | NP |
| C4956 | 92 | F | 23.3 | Ctrl | x | x | NP |
| PDC111 | 88 | F | 23 | Ctrl | x | x | NP |
| C090 | 83 | M | 21 | Ctrl | x | x | NP |
| PDC86 | 89 | M | 18 | Ctrl | x | x | NP |
| C077 | 84 | F | 23 | Ctrl | x | x | NP |
| C4340 | 47 | M | 13 | Ctrl |  | x | NP |
| C5610 | 50 | F | 11 | Ctrl |  | x | NP |
| PD596 | 85 | M | 16 | PD | x |  | 4 |
| PD204 | 86 | F | 5 | PD | x | x | 3 |
| PD745 | 81 | F | 20 | PD | x | x | 6 |
| PD759 | 83 | F | 15 | PD | x | x | 6 |
| PD831 | 77 | M | 15 | PD | x |  | 6 |
| PD154 | 79 | M | 19 | PD | x | x | 6 |
| PD788 | 78 | M | 9 | PD | x | x | 4 |
| PD721 | 82 | F | 24 | PD | x | x | 6 |
| PD3873 | 76 | F | 7 | PD | x |  | NP |
| PD816 | 80 | F | 20 | PD | x | x | 6 |
| PD801 | 90 | F | 24 | PD |  | x | 3 |
| PD955 | 85 | M | 12 | PD |  | x | 5 |
| PD917 | 77 | M | 24 | PD |  | x | 4 |

**Extended Data Table 4:**
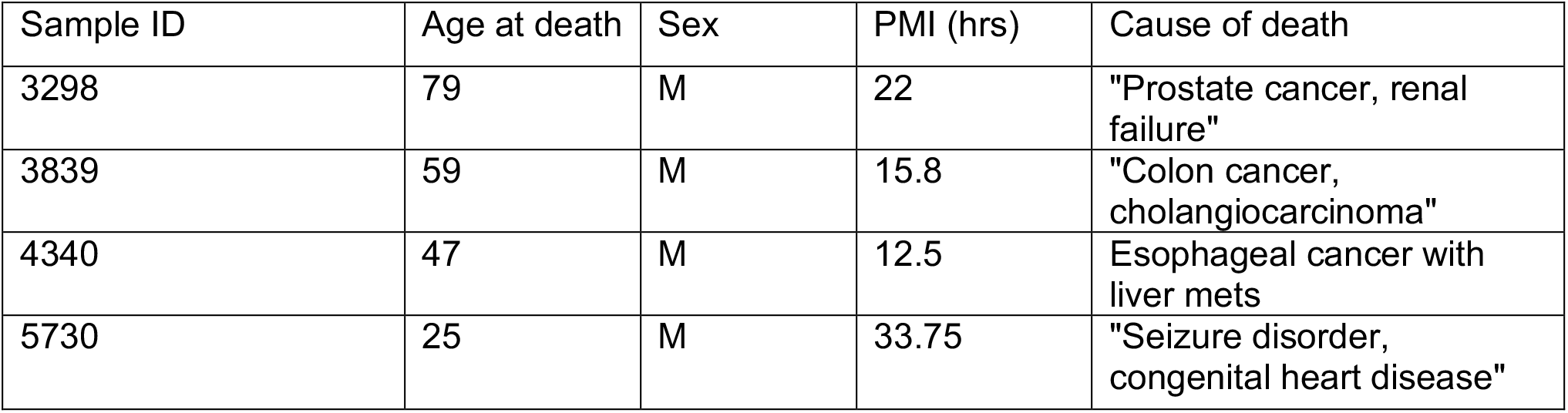
Basic demographics for 4 neuropathologically normal individuals for human caudate samples used for snRNA-seq experiments

| Sample ID | Age at death | Sex | PMI (hrs) | Cause of death |
| --- | --- | --- | --- | --- |
| 3298 | 79 | M | 22 | "Prostate cancer, renal failure" |
| 3839 | 59 | M | 15.8 | "Colon cancer, cholangiocarcinoma" |
| 4340 | 47 | M | 12.5 | Esophageal cancer with liver mets |
| 5730 | 25 | M | 33.75 | "Seizure disorder, congenital heart disease" |

